# Deep Neural Network Models of Object Recognition Exhibit Human-Like Limitations when Performing Visual Search Tasks

**DOI:** 10.1101/2020.10.26.354258

**Authors:** David A. Nicholson, Astrid A. Prinz

## Abstract

To find an object we are looking for, we must recognize it. Prevailing models of visual search neglect recognition, focusing instead on selective attention mechanisms. These models account for performance limitations that participants exhibit when searching highly simplified stimuli often used in laboratory tasks. However, it is unclear how to apply these models to complex natural images of real-world objects. Deep neural networks (DNN) can be applied to any image, and recently have emerged as state-of-the-art models of object recognition in the primate ventral visual pathway. Using these DNN models, we ask whether object recognition explains limitations on performance across visual search tasks. First, we show that DNNs exhibit a hallmark effect seen when participants search simplified stimuli. Further experiments show this effect results from optimizing for object recognition: DNNs trained from randomly-initialized weights do not exhibit the same performance limitations. Next, we test DNN models of object recognition with natural images, using a dataset where each image has a visual search difficulty score, derived from human reaction times. We find DNN accuracy is inversely correlated with visual search difficulty score. Our findings suggest that to a large extent visual search performance is explained by object recognition.

## INTRODUCTION

Visual search and object recognition are two real-world behaviors we engage in constantly, for example when we look for our keys on a cluttered desktop (Eckstein, 2011; Wolfe & Horowitz, 2017). Understanding how the brain solves the problems of object recognition and visual search is important for psychology, neuroscience, and artificial intelligence(DiCarlo et al., 2012; DiCarlo & Cox, 2007; Eckstein, 2011; Geisler & Cormack, 2011; Lindsay, 2020; Peelen & Kastner, 2014; Wolfe & Horowitz, 2017). Both behaviors have long been studied separately, but the two are obviously related: to find an object we are looking for, we must recognize it. As reviewed briefly below, previous studies point to a role for object recognition in visual search, but little computational work has been done to ask how a single model might account for both behaviors.

The dominant models of visual search neglect object recognition, instead focusing on selective attention mechanisms (Eckstein, 1998; Eckstein et al., 2000; Eckstein, 2011; J. Palmer et al., 2000; Treisman & Gelade, 1980; Wolfe et al., 1989; Wolfe, 1994; Wolfe & Gray, 2007). Selective attention mechanisms were proposed to account for performance limitations exhibited by participants in laboratory visual search tasks (Eckstein, 2011; Poder, 2017; Wolfe & Horowitz, 2017). These performance limitations are observed when participants search for targets in highly simplified stimuli, like those shown in Figure 1, that have been used in hundreds of studies (Wolfe, 1998). As can be seen in Figure 1, the stimuli typically consist of a two-dimensional array of items. Participants search for a target item that is distinguished from distractor items by one or two parametrically-defined features, such as hue, luminance, or orientation. Accordingly, the dominant models reproduce visual search behavior when given features and/or some number of items as inputs. We emphasize that, in spite of any differences, all prevailing models have this key characteristic in common: they are all specified (Cooper & Guest, 2014) such that they operate on sets of discrete items, where each item is described by only a few human-defined features. The well-known Guided Search model states this assumption about inputs explicitly (see p.104 of (Wolfe & Gray, 2007), “Another major simplification…”, and (Wolfe & Bennett, 1997)). Similarly, another widely-used family of visual search models (Eckstein, 1998; Eckstein et al., 2000; Eckstein, 2011; J. Palmer, 1994; J. Palmer et al., 2000) built within a signal detection theory framework (Green et al., 1966) typically represents each set of items as a distribution in a very low dimensional “feature space”, where each dimension of that space corresponds to a human-defined feature (Eckstein et al., 2000).

**Figure 1.**
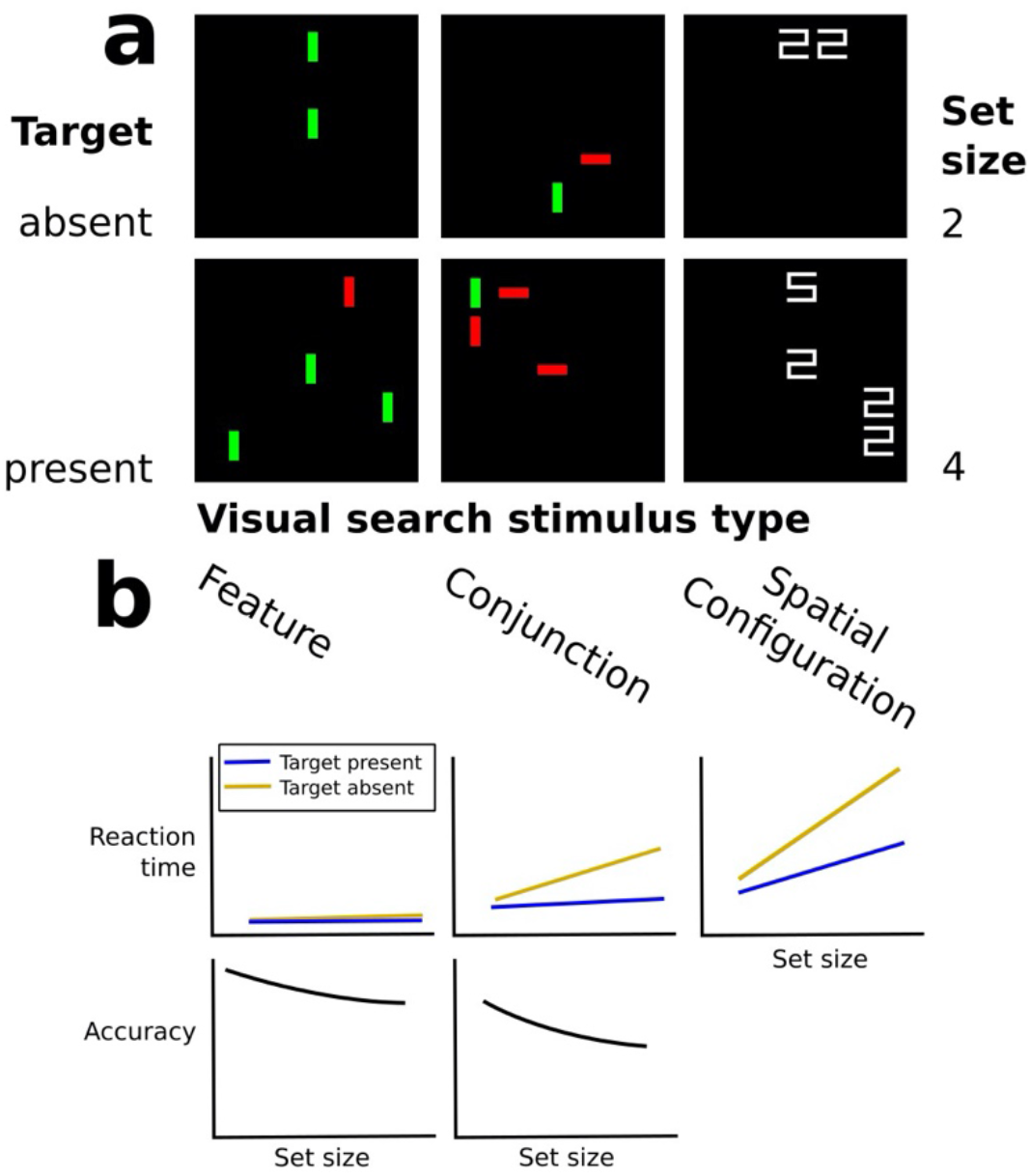
Set size effects are a hallmark finding from laboratory visual search tasks. Panel **a** shows example simplified displays commonly used in visual search tasks. In top row of **a** a target is absent and in the bottom row it is present. Displays in each row also have different set sizes (total number of items including target and distractors): on the top row of **a** the set size is two and the bottom row it is four. Panel **b** schematically depicts *set size effects*, redrawn from (Wolfe et al., 2010) and (Eckstein, 1998)). Effect size varies based on the features that distinguish targets from distractors (shown in columns). In the left column of **a**, the target can be distinguished from distractors by a single *feature*, color; in the middle column, by a *conjunction* of features, color and orientation; in the right column, by a *spatial configuration* of multiple features.

Because of how the prevailing models are specified, it remains unclear how to extend them to account for search for real-world objects in natural scenes, like those shown in Figure 6. In these scenes, an object we are looking for is not guaranteed to be distinguished from other objects by just one or two parametrically-defined features (Peelen & Kastner, 2014; Wolfe et al., 2011). In addition, natural scenes are not easily divided into discrete items (Hulleman & Olivers, 2017a, 2017b). Even if it were possible to easily extend existing models to natural images, their predictions might not match real world visual search behavior. For example, the key mechanism proposed by the Guided Search model is a serial processing step (Wolfe, 1994; Wolfe et al., 1989; Wolfe & Gray, 2007). If search involving multiple features were limited by serial processing, it would imply that cluttered natural scenes should produce large reaction times, but often searches of these scenes can be highly efficient (Kristjánsson, 2015; Nakayama & Martini, 2011; Peelen & Kastner, 2014; Wolfe et al., 2011). Although search of natural scenes can be remarkably efficient, participants do exhibit performance limitations. For example, some images produce much larger reaction times than others, as demonstrated experimentally, and features of natural scenes are predictive of reaction times (Ionescu et al., 2016; Katti et al., 2017).

**Figure 2.**
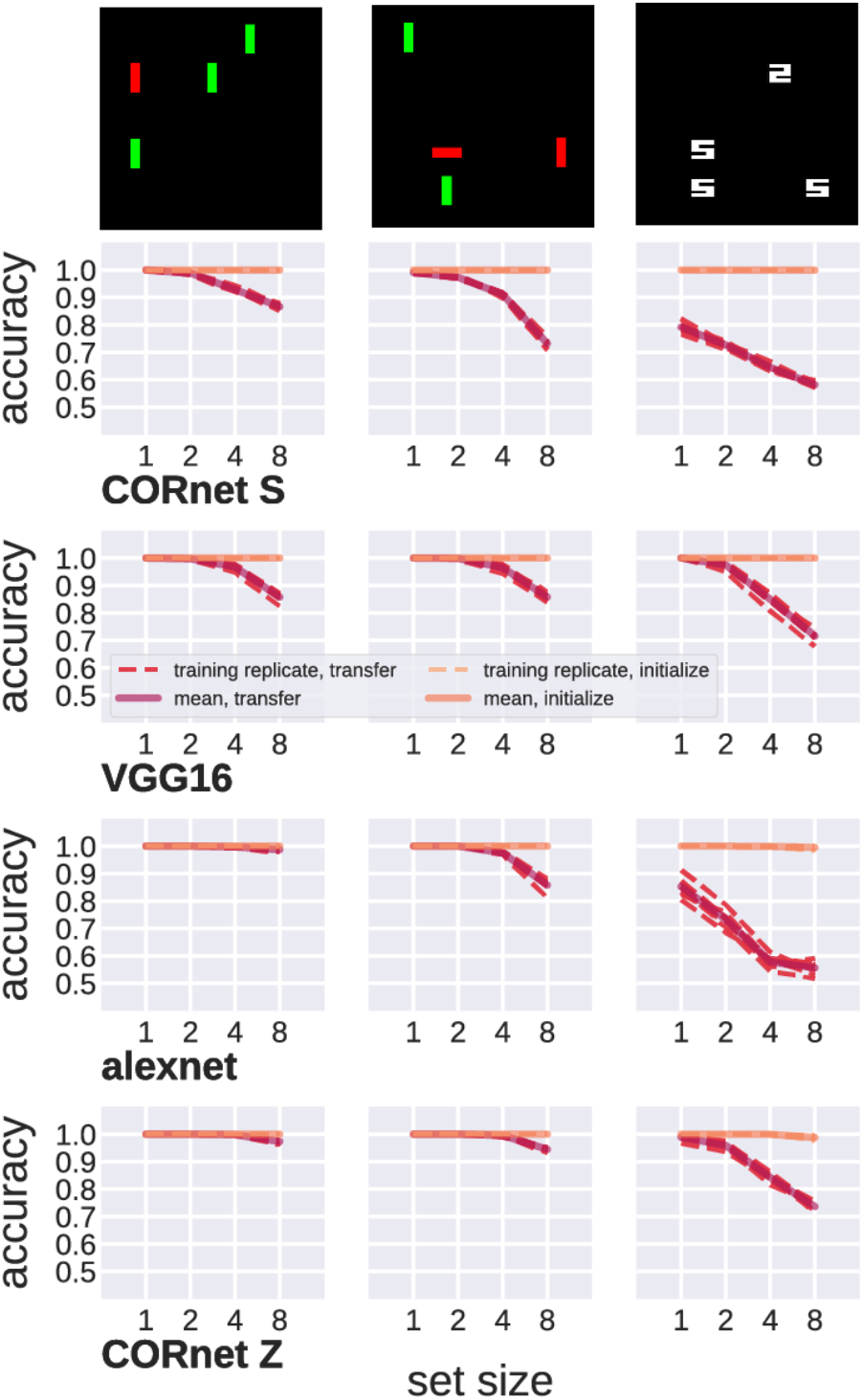
Accuracy as a function of set size, for DNN architectures trained to perform a visual search task. Rows show four different models trained on datasets consisting of 3 different search stimulus types (examples shown at top of columns). Each panel shows accuracy as a function of set size, for eight training replicates (dashed lines). Solid lines indicate mean across all trials and replicates. Brown dashed lines and red solid lines indicate accuracy for DNN-based object recognition models, that had weights pre-trained for image classification and were adapted to this visual search task using transfer learning. Peach dashed lines and salmon solid lines indicate accuracy for the same DNN architectures trained from randomly-initialized weights, not pre-trained for image classification. Columns are ordered by effect size, the difference in accuracy between set size 1 and 8 for the object recognition models. Rows are organized in increasing order of per-model effect size, averaged across all stimulus types (e.g., CORnet Z showed the smallest effect size when averaged across stimulus types). All example images for different stimulus types are shown with the target present condition. Stimulus types from left to right are: red vertical line target v. green vertical line distractors; red vertical line target v. red horizontal and green vertical line distractors; white digital 2 target v. white digital 5 distractors.

**Figure 3.**
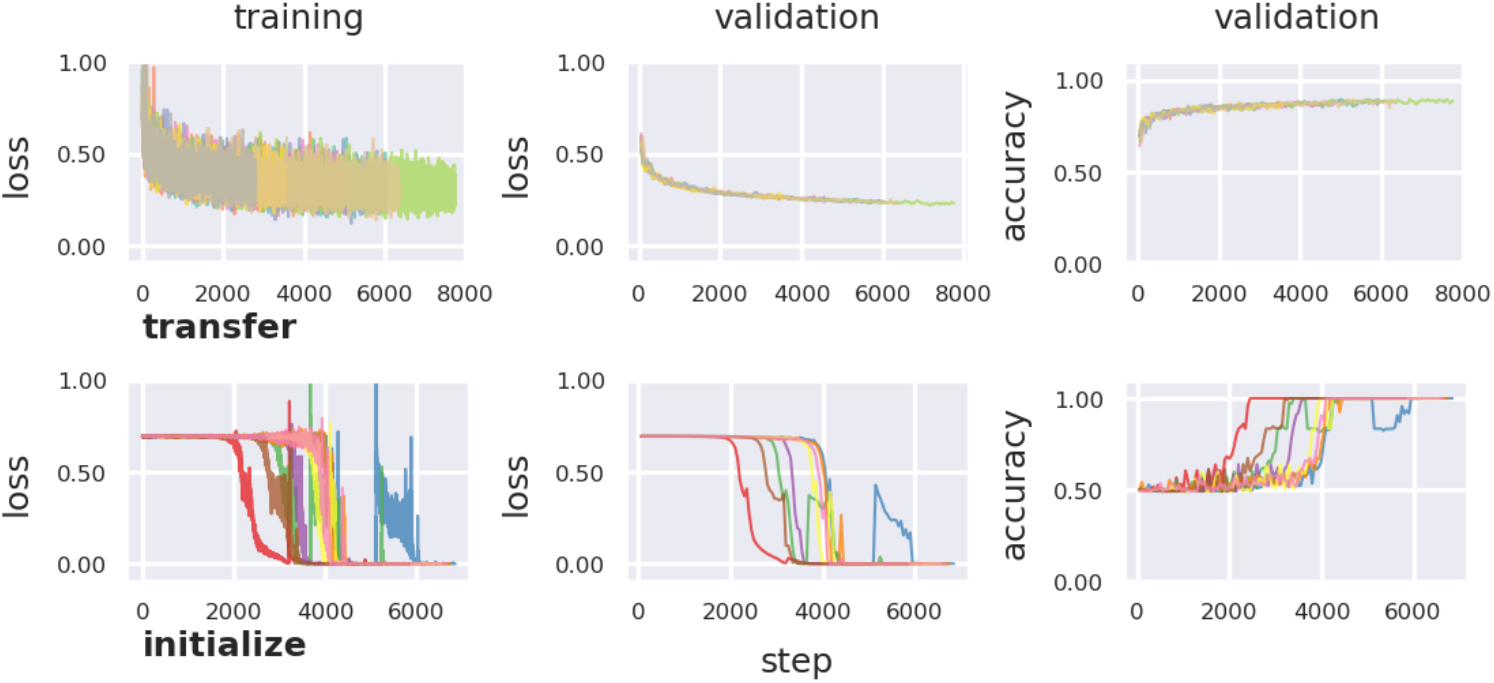
Training histories for DNN architecture trained on visual search task with simplified stimuli. Shown are representative training histories from the Alexnet model. (Plots of training histories for all models can be found in the openly shared repository accompanying this paper; see methods for link.) Top row, training histories from experiments using transfer learning to hold parameters fixed that were pre-trained for object recognition (indicated by “transfer”); bottom row, histories for the same model trained with randomly-initialized weights (indicated by “initialize”). For each DNN architecture and training method (transfer or from randomly initialization), we trained eight replicates, each indicated by a different hue. Each plot shows different metrics measured at different “steps” of training (a “step” is one batch of training data). First column shows the loss function used to optimize the model. Second column shows loss measured intermittently on a validation data set (not used to update model weights). The third column shows accuracy measured on the same validation set.

**Figure 4.**
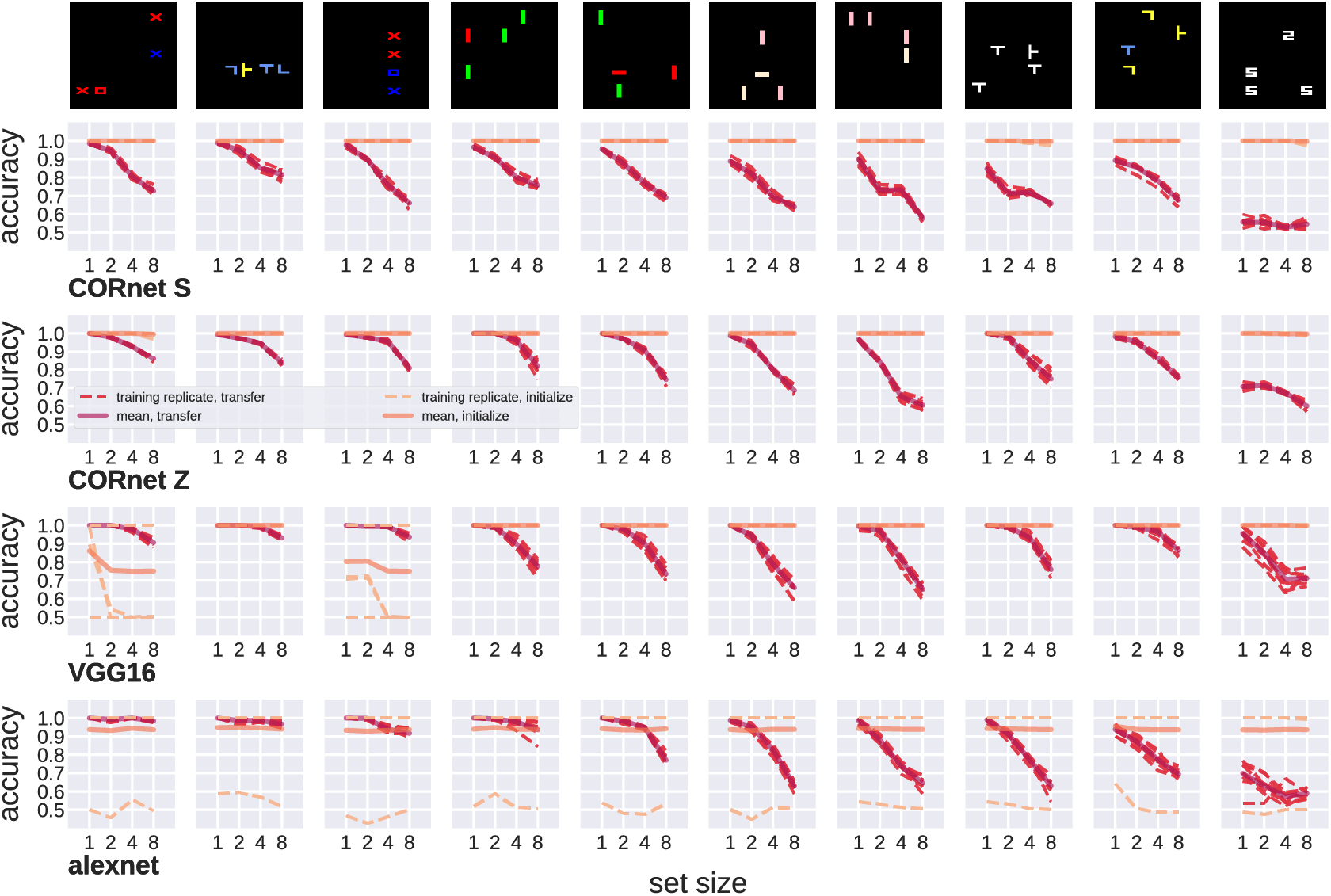
Accuracy as a function of set size, for DNN architectures trained to perform a visual search task. Here the dataset consisted of ten stimulus types, to test whether set size effects resulted from training DNNs to assign only one of two classes to many different stimulus types. Otherwise, plots are as in Figure 3. Stimulus types from left to right are: blue x target v. red x and o distractors; yellow rotated T target vs. blue T and L distractors; blue x target vs red x and blue o distractors; red vertical line target v. green vertical line distractors; red vertical line target v. red horizontal and green vertical line distractors; papaya-whip vertical line target v. cyan vertical line distractors; papapya-whip vertical line target v. papapya-whip horizontal and cyan vertical line distractors; rotated T target vs. T distractors; yellow T target vs. blue T and yellow L distractors; white digital 2 target v. white digital 5 distractors.

**Figure 5.**
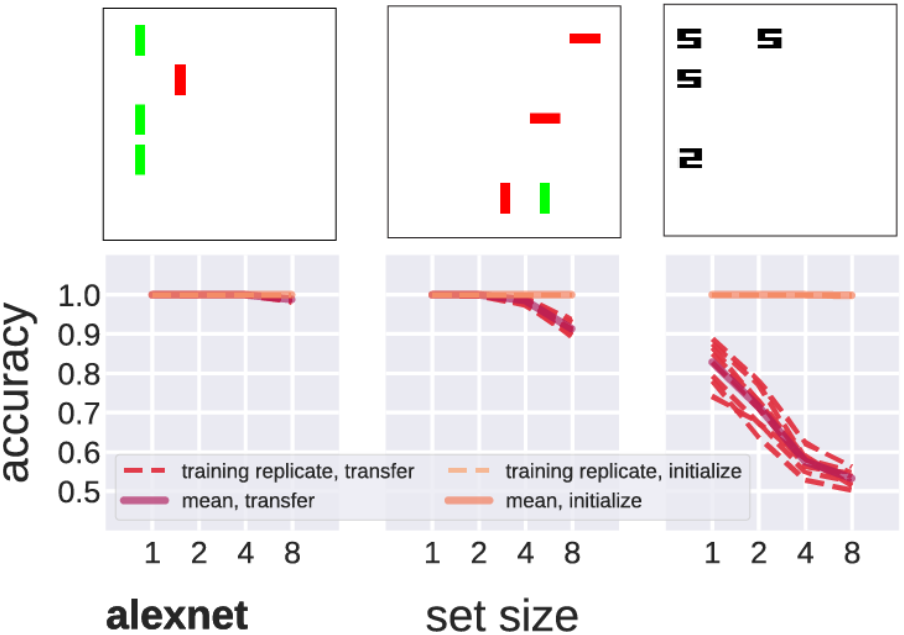
Accuracy as a function of set size, for DNN architectures trained to perform a visual search task. Here the dataset consisted of stimuli with white backgrounds, as a control experiment (details in text). Otherwise, plots are as in Figure 3.

**Figure 6.**
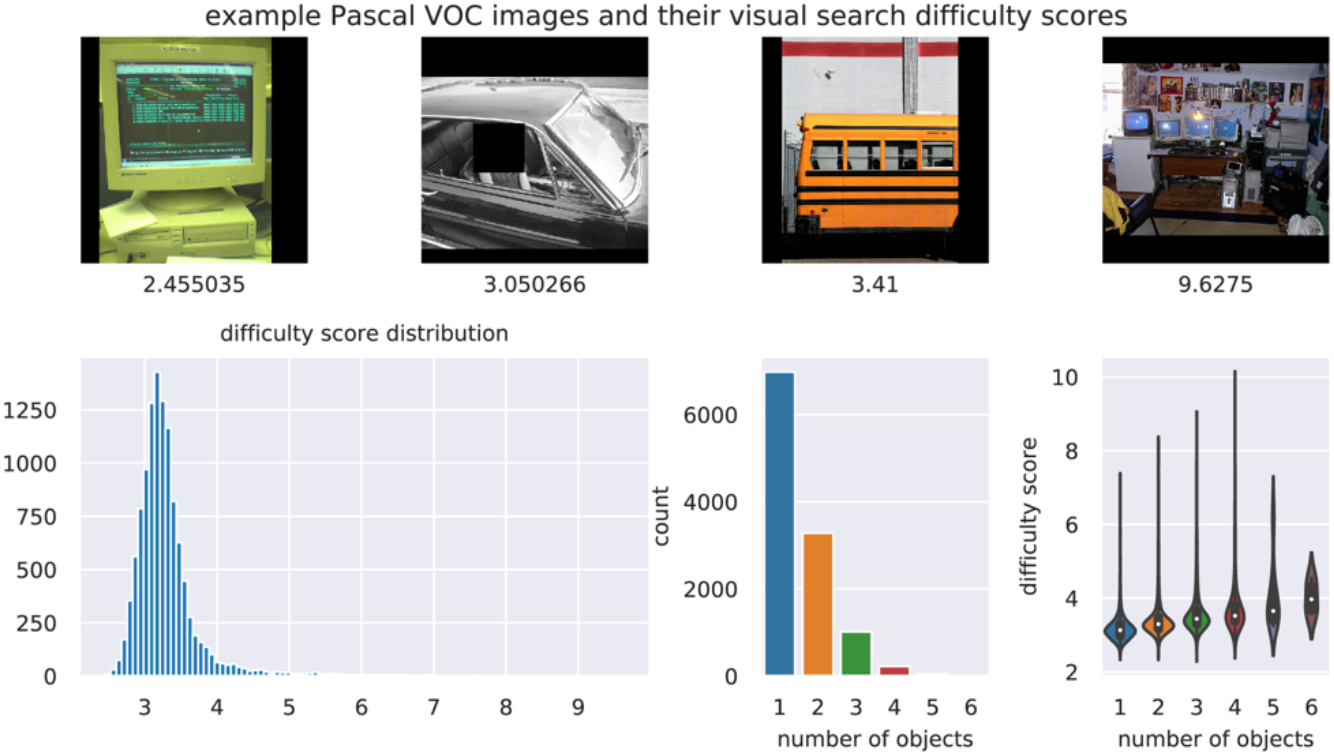
The Visual Search Difficulty dataset assigns search difficulty scores to images from the Pascal VOC dataset, based on reaction times of human subjects searching each image for targets. In the top row, example images from the Pascal VOC dataset are shown, and the visual search difficulty score assigned to that image is shown underneath it. Images are the first through fourth quartile of the distribution. Face in second image obscured for privacy. The distribution of scores is shown in the first panel of the bottom row. The middle panel in the bottom row shows the counts for the number of objects annotated per image. Last panel, bottom row shows distribution of difficulty scores for images, categorized by the number of objects annotated.

It might be the case that the limitations participants exhibit when searching natural images are explained in part by their ability to recognize the objects they are looking for. In experiments using simplified stimuli like those discussed above, performance limitations are explained in large part by target-distractor similarity (Bauer et al., 1996; Duncan & Humphreys, 1989; D’Zmura, 1991). In other words, the discriminability of the targets and distractors impairs performance. The importance of discriminability strongly suggests a direct link between visual search and object recognition, as discussed by (Nakayama & Martini, 2011). Models of object recognition in the visual system are often described in terms of discriminative machine learning models (DiCarlo & Cox, 2007; Nakayama & Martini, 2011), that classify inputs using learned decision boundaries. More recent work further implicates object recognition in visual search, showing that reaction times during search for real-world objects can be predicted by the similarity of neural representations in high-level cortical areas known to be responsible for object recognition (Cohen et al., 2017).

To ask how visual search relates to object recognition, and to address issues with dominant models of visual search, we turned to deep neural network (DNN) models. DNNs offer a key advantage compared to prevailing visual search models: DNNs are image-computable (Geisler & Cormack, 2011; D. L. Yamins et al., 2014a). This means that DNN models accept any image as input, whereas the dominant visual search models are limited to images from which a human-specified feature representation can be extracted, as just described. If applied to visual search tasks, DNNs could provide predictions both for the highly simplified stimuli used in lab tasks and for relatively complex natural scenes. Recently, the study of object recognition has been revolutionized by the use of DNN models. When optimized for task performance, using methods that have come to be called deep learning (Richards et al., 2019; D. L. Yamins et al., 2014b; D. L. K. Yamins & DiCarlo, 2016), DNN models accurately predict both neural activity in the primate ventral visual pathway and behavior during object recognition tasks (Schrimpf et al., 2018; D. L. K. Yamins & DiCarlo, 2016). Specifically, DNN accuracy is correlated with accuracy of human subjects and nonhuman primates performing object recognition tasks. Therefore, to better understand the relationship between visual search and object recognition, we utilize DNNs that are currently best-in-class models of object recognition.

We hypothesize that performance limitations across stimuli used for visual search tasks are constrained in part by object recognition. A strength of the deep learning framework for neuroscience is that it allows modelers to ask how the brain being optimized for one task might impact how it performs other tasks (Kell & McDermott, 2019). This approach is analogous to ideal observer models (Geisler, 2003; Geisler & Cormack, 2011) which have been applied successfully to visual search. Ideal observer models provide insights by adopting a normative approach: proposing a closed-form optimal solution for tasks, and then asking how real-world behavior deviates from the behavior dictated by the optimal solution. DNN models obviously do not provide a closed-formed solution for tasks. However, as noted previously (Kell & McDermott, 2019), the optimization perspective has yielded a significant body of empirical evidence that DNN models perform “near ideally” (at least as measured with a test data set, that models do not see during training). In this way, we use a normative approach to theoretically link object recognition and visual search, even though we cannot derive a closed-form solution for many object recognition and visual search tasks that use complex natural images.

Thus, we chose the deep learning framework to test whether DNNs optimized for object recognition exhibit human-like limitations on performance across the different types of images used in visual search tasks: the highly-simplified stimuli used for controlled experiments, and relatively complex natural scenes containing real objects. Our focus on object recognition as a constraint on visual search is distinct from previous studies that have employed various neural network models to investigate visual search (Eckstein, 2011; Grossberg et al., 1994; Ma et al., 2011; Poder, 2017) and selective visual attention (Bobier et al., 2014; Lindsay, 2020; Lindsay & Miller, 2017). To measure the behavior of DNN-based object recognition models performing visual search tasks, we use methods from deep learning to adapt pre-trained models to new tasks. As we show below, this approach produced results consistent with the idea that object recognition places constraints on visual search performance.

## METHODS

### NEURAL NETWORK ARCHITECTURES

We chose four neural network architectures as a representative sample of DNN-based object recognition models, to increase the likelihood that our results are general and not an artifact of any specific architecture. All the models we test are convolutional neural networks (Krizhevsky et al., 2012) where the nonlinearity applied after each layer is the rectifier activation function (Glorot et al., 2011). Two of the models, AlexNet (Krizhevsky et al., 2012) and VGG16 (Simonyan & Zisserman, 2014) represented key advances in image classification by the computer vision community, and were later used in some of the first papers that proposed DNNs as models of object recognition (Cadieu et al., 2014; Jozwik et al., 2017). The other two, CORnet S and CORnet Z, are two DNNs specifically developed in pursuit of good performance under a metric proposed to account for a model’s ability to predict both brain activity and behavior during object recognition tasks (Schrimpf et al., 2018).

### TRANSFER LEARNING

For all experiments, we employ transfer learning methods used with deep neural network models (Kornblith et al., 2019; Yosinski et al., 2014). Essentially, we hold fixed all parameters optimized for object recognition, except for those in the final output layer of the neural network. We replace the final layer used for image classification with a new layer that has an appropriate number of units for the visual search task, and then adapt the model to this task by optimizing for performance with a training set. For standard lab tasks using simplified stimuli, the final layer has two output units corresponding to “target present” and “target absent”; for tasks using natural scenes, the number of output units corresponds to the number of classes in the dataset (20 in the case of the Pascal VOC dataset we use).

To assess performance during and after transfer learning, we followed good practices for machine learning (Hastie et al., 2001). These included dividing datasets into training, validation, and test subsets, where the validation set was used to evaluate the model during training, and the test set was withheld during training and used to measure model behavior afterward. For standard lab tasks using simplified stimuli, all DNN models were trained on a dataset consisting of all different types of stimuli (see for example columns in Figure 2 and Figure 4), with 1200 samples for each type. During training, batches were drawn randomly from this dataset, without regard for stimulus type. The results we report in Figure 2 used three types, and the results in Figure 4 used ten types. Stimuli were generated with jitter in the placement of the items, to guarantee that there were no repeated images that would encourage the DNNs to simply memorize the correct answer during training. 1200 samples was the maximum number we could generate per stimulus type without repeats, given the parameters we used to create them. For tasks using natural scenes from the Pascal VOC 2012 dataset (Everingham et al., 2012), we split the data as was done in the original study (Ionescu et al., 2016) that assigned visual search difficulty scores to the Pascal VOC dataset, since we use those scores. That is, we used 50% of the Pascal VOC training-validation sets as our training set, 25% as a validation set, and 25% as a test set.

### TRAINING METHODS AND ANALYSIS FOR VISUAL SEARCH DIFFICULTY DATASET

In Experiment 2 we compute different measures of accuracy for DNN models trained on the Pascal VOC dataset. This allows us to ask whether those measures correlate with visual search difficulty scores derived from reaction times of human participants searching Pascal VOC images for targets.

As stated above, we used the same transfer learning method for this experiment that we used for Experiment 1. This allowed us to directly compare model behavior between experiments. However, we needed to extend our approach to deal with an important difference between the datasets used in the two experiments: in the Pascal VOC dataset, multiple targets can be present. To minimize the possibility that our results depended on how we trained models, we tested multiple methods of training. In addition to the standard single-label image classification task, in which DNNs are trained using cross entropy as the loss function, we trained models for multi-label image classification, where models are trained using a binary cross-entropy loss. In the latter setting, the vector of predictions that a DNN returns is interpreted as independent probabilities that each label, i.e., class, can be applied to the image. Regardless of whether we were training models for single-label or multi-label classification, we used all images, to avoid minimizing the amount of training data. When training for single-label classification, then, we chose a single label for images with multiple labels, using one of two methods: (1) pick the label corresponding to the largest object in the image, as measured by the area of its annotated bounding box, or (2) pick a random object. As we show in the results, testing both approaches demonstrated that which object we chose had a negligible effect on results (accuracies as measured on the test set were 1-2% lower for randomly chosen labels, but this had no impact on the correlations we report between accuracy and visual search difficulty scores). For multi-label classification, our accuracy measure was “macro” F1 score as implemented in scikit-learn (Grisel et al., 2020; Pedregosa et al., 2011), which averages accuracy across classes.

To analyze the behavior of trained models, we treated their predictions as “trials”, where the model reported on each trial whether a class was present or absent. We considered each element of the vector of predictions produced by a model as one “trial”. We did this regardless of whether the model was trained for single-label or multi-label classification. Thus, we treated every model trained on Pascal VOC as if for each image it performed 20 trials of reporting target present or absent, since there are 20 classes labeled in the dataset. However, when models were trained for single-label classification, we only considered predictions for images in the test set with a single label. This means accuracies we reported were not lowered artificially because of images with multiple labels. We point out that a significant portion of images in Pascal VOC have just a single label, as shown in Figure 6. For models trained for multi-label classification, we considered all images regardless of the number of labels. We then used these trials to compute accuracies shown in Figure 7, as we now explain.

**Figure 7.**
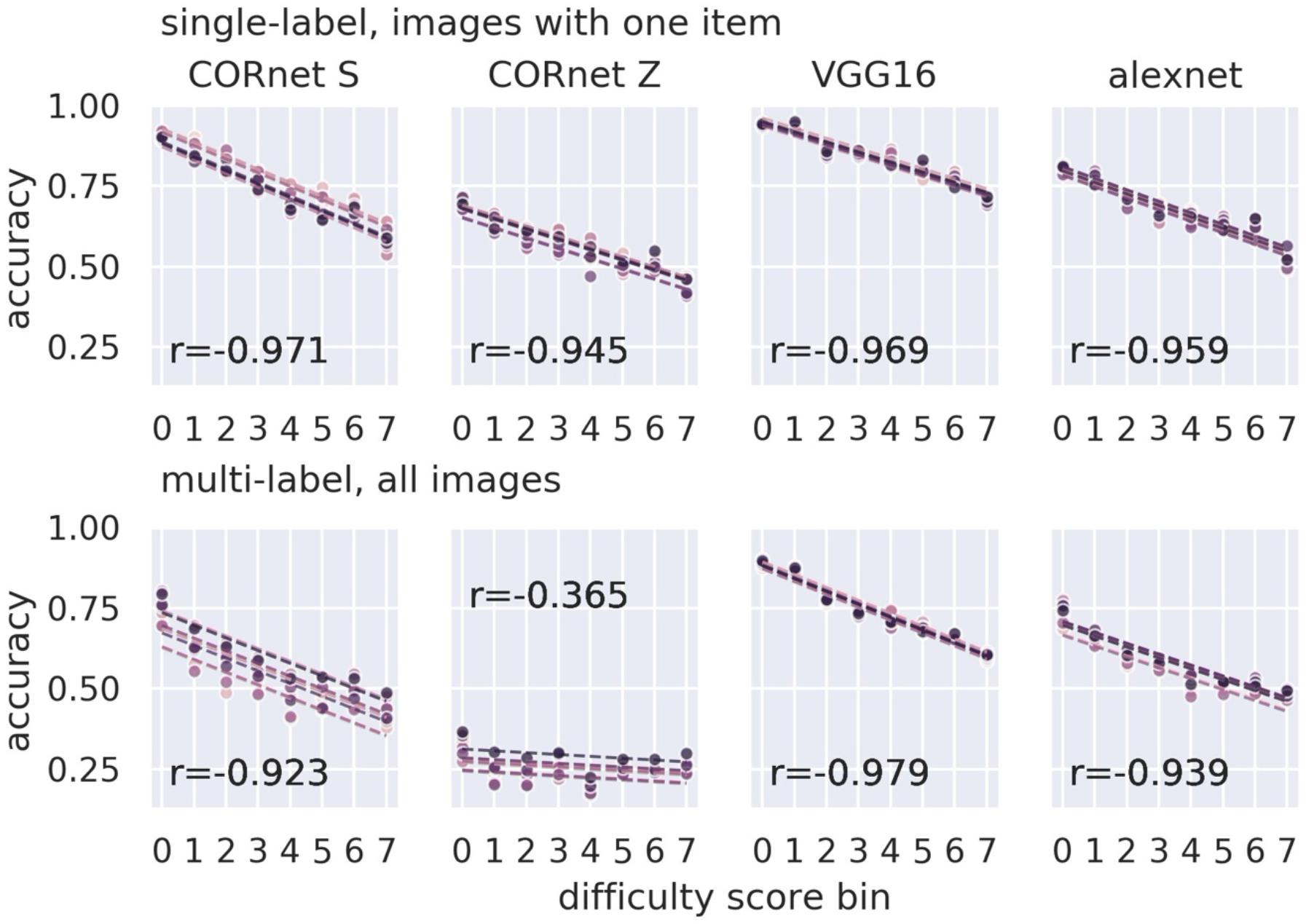
Correlation of target detection accuracy of DNN models of object recognition and visual search difficulty scores derived from reaction times of human participants. Plots show clear correlation between binned visual search difficulty scores (x axes) and accuracy of DNN-based object recognition models when measured on trials within a given bin (y axes). Dots indicate accuracy within each bin of model reporting target class present or absent on each trial as described in methods. Different colors indicate training replicate. Dashed line shows a model fit using linear correlation. R value is for repeated measures correlation. Different models are shown in columns, with single-label (top row) or multi-label (bottom row) classification.

To ask whether there was a relationship between model accuracy and behavior of human participants performing visual search tasks with the same images, we binned the Visual Search Difficulty scores assigned to the Pascal VOC images based on human reaction times. We then computed model accuracy within these bins using the trials as just described. We binned such that there were equal numbers of trials in each bin. Note that accuracies are only measured for a set of images within a bin *after* the images have been binned by difficulty scores. That is, the accuracy values do not result from taking a mean of multiple accuracies within any given bin. We also varied the number of bins, and found that this did not qualitatively change our key result. We also tested another binning scheme, using bins of equal width, but this produced bins with very few images, because of the uneven distribution of visual difficulty scores, as shown in Figure 6. Even when using as few as four bins with equal widths, we still had bins with a few as 8 images, because of the long tail of the difficulty score distribution. In contrast, we typically had 1000s of images per bin when using the strategy that divided up bins so that they contained equal numbers of images. Thus, we adopted the latter binning strategy, because our main goal was to compute a good estimate of accuracy within each bin, and we had no compelling reasons to use other binning schemes.

After binning the difficulty scores and computing trial accuracies within those bins, we tested for a relationship with a repeated measures correlation (Bakdash & Marusich, 2017), using the implementation in the pingouin library (Vallat, 2018). This statistical technique tests for a within-individual correlation between paired measures (accuracy and visual search difficulty score) when measured across individuals on multiple occasions. We treated each of the eight training replicates as if they were individuals, and each of the bins as separate, possibly non-independent, measures, and then computed the repeated measures correlation accordingly.

### CODE AVAILABILITY

To aid with reproducibility of our experiments, and to make them more accessible to other researchers, we developed a separate software library, `visual-search-nets`, available at https://github.com/NickleDave/visual-search-nets. We also developed a tool to generate datasets of the simplified visual search stimuli like those we use in Experiment 1, in a format that is convenient for training neural networks, available at https://github.com/NickleDave/searchstims. All configuration files for carrying out experiments, and scripts for generating stimuli, analyzing results and creating figures, are available in the repository corresponding to this manuscript: https://github.com/NickleDave/Nicholson-Prinz-2020. Libraries, tools, and code for analysis and figures were developed with the following Python libraries: attrs (Schlawack, 2019), numpy (Harris et al., 2020; Walt et al., 2011), scipy (Virtanen et al., 2019), scikit-learn (Grisel et al., 2020; Pedregosa et al., 2011), pandas (team, 2020; Wes McKinney, 2010), matplotlib (Caswell et al., 2020; Hunter, 2007), seaborn (Waskom et al., 2020), jupyter (Kluyver et al., 2016), pingouin (Vallat, 2018), pygame (Schinners, 2019), pytorch (Paszke et al., 2019), statsmodels (Seabold & Perktold, 2010), and torchvision (Marcel & Rodriguez, 2010).

## RESULTS

In order to test whether DNN-based object detection models exhibit human-like limitations when performing visual search tasks, we employ transfer learning methods used with deep neural network models (Kornblith et al., 2019; Yosinski et al., 2014) (see Transfer learning in Methods). Essentially, we hold fixed all neural network parameters optimized for object recognition, except for those in the final output layer. We replace the final layer used for image classification with a new layer that has an appropriate number of units for the visual search task. Then we divide datasets into training, validation, and test subsets, using the validation set to evaluate the model during training, and using the held-out test set to measure model behavior with data not seeing during training. We measure the behavior of four DNN architectures used as models of object recognition in the primate ventral visual pathway: AlexNet (Krizhevsky et al., 2012), VGG16 (Simonyan & Zisserman, 2014) used in (Cadieu et al., 2014; Jozwik et al., 2017), and CORnet S and CORnet Z (Schrimpf et al., 2018). We chose these four architectures as a representative sample of DNN-based object recognition models, to increase the likelihood that our results are not an artifact of any specific architecture.

### EXPERIMENT 1

#### SET SIZE EFFECTS ARE A HALLMARK OF SEARCH TASKS THAT USE SIMPLIFIED STIMULI

We first tested whether DNNs exhibited behavior similar to that of humans performing laboratory visual search tasks carried out with highly-simplified stimuli like those shown in in Figure 1. The prevailing models of visual search are built on results obtained with these stimuli, which have been employed in hundreds of studies. As referenced in the introduction, there are essentially two competing families of models (Eckstein, 2011; Wolfe, n.d., 1994). The evidence for both consists of what are called “set size effects”. Accordingly, we test whether DNN-based object recognition models also exhibit set size effects. To define what set size effects are, we briefly describe how these laboratory search tasks are typically performed (Wolfe, 1998). On each trial, the participant views one of the highly simplified stimuli and reports whether a target is present (Figure 1**a** bottom row) or absent (Figure 1**a** top row) among distractors. The term “set size” refers to the total number of items, i.e., distractors plus target when present. By extension, the term “set size effect” refers to any change in some behavioral measure of target detection, such as reaction time or accuracy, that depends on increasing the set size, usually by increasing the number of distractors. Schematic depictions of results that would indicate set size effects are shown in Figure 1**b**. These set size effects are taken as evidence for different types of computations thought to be involved in visual search. Therefore, we tested whether DNN models of object recognition exhibit set size effects, supporting the idea that optimizing for object recognition impacts visual search performance. Specifically, we asked whether DNNs showed a decrease in accuracy of detecting a target as the number of distractors increased. We chose accuracy as a measure because it is easily computed for standard DNN models, and has typically been used as a behavioral measure in studies of object detection utilizing DNN models. We address other behavioral measures such as reaction times in the discussion.

#### DNNS PRE-TRAINED FOR OBJECT RECOGNITION SHOW SET SIZE EFFECTS, WHILE DNNS TRAINED FROM RANDOMLY INITIALIZED WEIGHTS ACHIEVE PERFECT ACCURACY

Our key results from this experiment, as shown in Figure 2, Figure 3, and Figure 4, are as follows: all DNN-based models of object recognition we tested exhibited set size effects; the ranking of the effect size across models was qualitatively the same as the ranking exhibited by human participants performing visual search tasks; and this effect resulted from optimizing the DNN architectures for object recognition.

We explain these results in detail. First, we used transfer learning to adapt four different DNN architectures that have been used as models of object recognition (rows in Figure 2) to the visual search task. We trained eight replicates of each DNN architecture on a dataset consisting of three types of visual search stimuli (columns in Figure 2) with 1200 samples for each type (see Methods for details). For clarity, we emphasize that all dataset splits (training, validation, and test) contained all three stimulus types. All models showed a drop in target detection accuracy as the number of distractors increased (solid purple line). This set size effect was consistent across the eight training replicates we ran for each model (dashed red lines). The effect size was different for different stimulus types; we measured effect size by taking the difference between accuracy for set size 8 and accuracy for set size 1. The columns in Figure 2 are sorted by the size of this effect, in increasing order, averaged across models. We observed that when sorted by set size effect, the stimulus types were ranked in the same way they would be by researchers working with the dominant models of visual search. I.e., stimuli thought to give rise to “efficient” search in humans were easily discriminable by the networks, like the red vertical target vs. green vertical distractors in the first column in Figure 2. Similarly, stimulus types thought to give rise to “inefficient” search, such as the “digital 2 target vs. digital 5 distractors stimulus in the third column, were difficult for the networks to discriminate. To underscore the strength of our approach, we emphasize that the latter stimulus in particular is an issue for prevailing models, because it is unclear how to break down the digital 2s and 5s into features (E. M. Palmer et al., 2011).

This initial finding demonstrated that DNN models exhibit set size effects, but it left unanswered the question of whether these effects are specific to object recognition models, or alternatively whether any DNN would exhibit similar set size effects. To answer this question, we carried out another set of experiments where we trained the exact same DNN architectures from randomly-initialized weights, instead of adapting object recognition models (with weights pre-trained for image classification on ImageNet). In all cases, DNNs trained from randomly-initialized weights achieve nearly perfect accuracy across all set sizes (solid salmon-colored lines in Figure 2). This effect held for all training replicates (dashed peach-colored lines). All neural network architectures tested achieved this level of accuracy.

To further illustrate this difference, we present in Table 1 the accuracy of all models we trained, measured across the entire test set (rather than measured per visual search stimulus set size). The results in the table make it clear that all DNN architectures trained from randomly-initialized weights achieved perfect or near perfect (~99%) accuracy on this task, while the exact same architectures achieved lower accuracy when weights were pre-trained for object recognition. Therefore, we conclude that the set size effects resulted from optimizing the models for object recognition, before using transfer learning to adapt them to the visual search task.

**Table 1.**
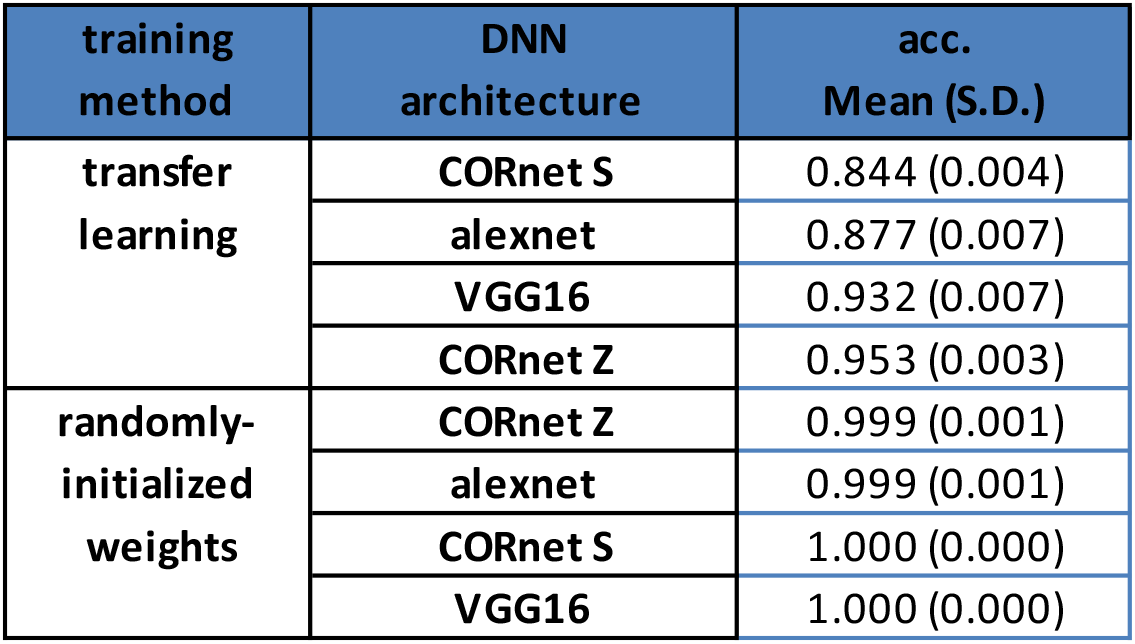

An alternative explanation for our findings would be that we simply failed to find the best method to adapt models using transfer learning. We emphasize that the method we chose, where we froze all weights pre-trained for object recognition before training a new final layer for visual search tasks, is meant to explicitly test whether object recognition models exhibited set size effects. Freezing all weights except for the last layer clearly limits our ability to improve models’ performance. However, even models trained with transfer learning performed quite well as measured on the test set, as shown in Table 1. We also eliminated the possibility that set size effects arose from other factors of our training method in preliminary experiments (Nicholson and Prinz, 2019). In those preliminary experiments, we examined the effect of hyperparameters such as learning rate, imbalance in the data set, and the size of the training set. Essentially, we increased the learning rate to achieve the maximum accuracy possible without overfitting, and we increased the dataset to the largest size possible without generating multiple examples of the same image. We also found in those preliminary experiments that balancing the dataset across visual search set sizes, as we did here, produced the best accuracy.

In addition, we took several steps to minimize the possibility that the results presented here were an artifact of our training method. Those steps included logging metrics at each step of training, then visually assessing plots of the logged training histories for evidence of overfitting to the training set, or failure of the optimization to converge. In Figure 3 we show representative training histories from the Alexnet model. (Plots of training histories for all models can be found in the code repository accompanying this paper; see Code availability section in Methods for link.) In almost all cases, we saw by plotting the loss values that the optimization converged (first column, Figure 3), and that loss also decreased when measured on a validation set that DNN models did not see during training (middle column, Figure 3). Convergence was smooth for models pre-trained on ImageNet where we only fit a new last layer (top row, Figure 3), compared to models trained from randomly initialized weights (bottom row, Figure 3), but in both cases the optimization clearly converged. We also saw that the models achieved high accuracy on the validation set (last column, Figure 3), indicating that what they learned during training generalized to unseen data. Hence, we do not find any evidence that set size effects can be attributed to an artifact of training.

Another potential concern with these findings is that the set size effects might be an artifact of training DNNs to assign multiple stimulus types to only one of two classes: target present or absent. However, in our preliminary studies (Nicholson & Prinz, 2019), we also saw set size effects when training DNNs on only a single stimulus type. To further test whether set size effects arose from mapping multiple stimulus types to just two classes, we ran experiments with ten stimulus types instead of three. Our results, shown in Figure 4, are qualitatively similar to those obtained with only three stimulus types. We saw that models were clearly able to effectively classify multiple stimulus types, yet the set size effect remained (solid red lines) By ranking stimulus types by effect size as before, we did observe that stimulus types that would be classically considered to be a “spatial configuration” of features, such as the digital 2 v. digital 5 stimulus, were more difficult for some models to classify when the dataset included more stimulus types (right 3 columns of Figure 4). When repeating the experiment where we trained the same DNN architectures from randomly-initialized weights, we again observed that for all architectures, most of the eight replicates achieved near perfect accuracies (dashed peach lines). The only exceptions were one replicate of AlexNet that failed to converge, and one replicate of VGG16 that did not achieve high accuracy on two of the ten stimulus types (third and fourth rows in Figure 4). These results serve as an “existence proof” that it is possible for these DNN models to find near-optimal solutions to this task, even when trained on ten stimuli. Finally, we saw by inspecting the training histories for all models that optimization converged even when training on these very large datasets (20.8k samples including training, validation and test sets), and models still obtained quite high accuracy on the test set (>85%).

A final concern that might be raised about our results is that the simplified stimuli might change the statistics of activations within the hidden layers of the neural networks in a way that impedes networks’ ability to learn the task. For example, the black backgrounds might produce lower activations on average than the activations produced by full-color images from ImageNet used when training models for object recognition. To address this concern, we carried out a control experiment where we produced the same set of simplified stimulus types, only with a white background instead of black, and we repeated the training with the AlexNet model. We again saw that AlexNet models pre-trained on ImageNet exhibited set size effects, whereas AlexNet models trained from randomly-initialized weights were able to achieve very high accuracy on the same task, as shown in Figure 5. Thus, this control experiment did not produce evidence that our result is an artifact of the stimulus.

To summarize, we conclude that DNN models of object recognition exhibit set size effects when performing visual search tasks that employ the standard highly-simplified stimuli used in the laboratory. We do not find these results can be explained as an artifact of training methods or the stimulus types used. Further experiments demonstrated that set size effects result from optimizing DNN architectures for object recognition; when training the same architectures from randomly-initialized weights, they were able to find solutions that allowed them perform the tasks with near perfect accuracy. Therefore, the set size effects we observed resulted from DNNs being optimized for object recognition before we used transfer learning to measure how they performed visual search tasks.

### EXPERIMENT 2

#### THE VISUAL SEARCH DIFFICULTY DATASET

As described in the introduction, a key limitation of dominant models of visual search is that it is unclear how to extend them to account for search of complex natural scenes containing real objects. A key finding from experiments using natural scenes to study visual search is that it can be highly efficient, as suggested by reaction times. In spite of this efficiency, different natural scenes elicit different reaction times, as evidenced by previous attempts to estimate the extent to which targets and contextual information influence real world visual search (Katti et al., 2017; Ionescu et al., 2016). Although the DNN models we test here do not directly produce reaction times as an output, we can ask whether their accuracy correlates with reaction times.

To test for a relationship between DNN accuracy and human reaction times during visual search, we use the Visual Search Difficulty dataset, first presented in (Ionescu et al., 2016). The dataset consists of search difficulty scores assigned to all images in Pascal VOC, a dataset developed to benchmark computer vision models. The difficulty scores are a weighted combination of reaction times of human participants asked to report whether a target is present or absent in the image (please see (Ionescu et al., 2016) for details). In Figure 6 we present representative images from Pascal VOC with their visual search difficulty scores (top row), as well as summary statistics of the Visual Search Difficulty dataset (bottom row). Briefly we note some key features. As shown in the first panel of the bottom row, the distribution of scores has a long tail: the median is around 3.25, but the range of scores is approximately 2-10. We also note that about half of the images have just a single object annotated, although nearly as many have two or three objects annotated, as shown in the middle panel of the bottom row. Lastly, we observed a slight trend where images with more objects had a higher difficulty score, as shown in the last panel in the bottom row, although we did not compute a correlation.

#### TARGET DETECTION ACCURACY OF DNN MODELS OF OBJECT RECOGNITION CORRELATES WITH VISUAL SEARCH DIFFICULTY SCORES

In Experiment 1 we were able to use the standard machine learning task that researchers studying object recognition use when adapting our models to perform visual search tasks: single-label image classification. In other words, we trained models to classify images with a single label, “target present” or “target absent”, and then treated the output as if the models were participants performing the laboratory visual search task. For Experiment 2, there are images with multiple annotated objects in this dataset, raising the question of what machine learning task we should use to adapt models to “perform” the laboratory task. (Unfortunately”task” is the term most frequently used in machine learning to relate a problem formulation to the objective function used when optimizing models, so we use it here even though we recognize it overloads the term we use to refer to behavioral experiments.) We tested three machine learning tasks, to show whether our results were specific to the task used, or a more general phenomenon. First, we adapted models using the same single-label image classification task we used for Experiment 1, but we did this in two ways. We trained models using either the largest object in an image, as defined by its bounding box, or a randomly chosen object. Results were essentially the same regardless of which object we chose, as indicated by model performance on the test set, as shown in Table 2. Note that a significant portion of the dataset contains images where only a single object is annotated, as we show in Figure 6. We then carried out a separate set of experiments where we trained models a third way: multi-label classification, meaning that the model could assign to a given image any number of labels corresponding to the twenty classes present in the dataset. To train models for multi-label classification, we used a standard method, adopting binary cross entropy as the loss function, and making the final output layer sigmoid units. This method essentially treats each label independently, as if we were applying binary classifiers for each class to the image. We then measured performance of these models with the f1 score, a metric commonly used for multi-label classification tasks (Li et al., 2017; Nam et al., 2014; Wang et al., 2016), that can be thought of as the weighted harmonic mean of precision and recall (Grisel et al., 2020). Specifically, we computed the “macro” f1 score, which averages the f1 score for each class.

**Table 2.**

| task<br>(machine learning) | DNN<br>architecture | acc.<br>(largest object) | acc.<br>(random object) | f1<br>(macro) |
| --- | --- | --- | --- | --- |
| single-label,<br>largest | CORnet Z | 0.542 (0.013) | 0.486 (0.009) | 0.217 (0.009) |
|  | alexnet | 0.652 (0.010) | 0.576 (0.009) | 0.226 (0.002) |
|  | CORnet S | 0.743 (0.013) | 0.656 (0.014) | 0.237 (0.009) |
|  | VGG16 | 0.786 (0.007) | 0.683 (0.007) | 0.242 (0.002) |
| single-label,<br>random | CORnet Z | 0.520 (0.016) | 0.501 (0.012) | 0.212 (0.022) |
|  | alexnet | 0.616 (0.008) | 0.587 (0.007) | 0.231 (0.001) |
|  | CORnet S | 0.666 (0.019) | 0.654 (0.015) | 0.243 (0.008) |
|  | VGG16 | 0.742 (0.009) | 0.709 (0.008) | 0.248 (0.002) |
| multi-label | CORnet Z | 0.406 (0.018) | 0.418 (0.016) | 0.113 (0.011) |
|  | alexnet | 0.614 (0.010) | 0.589 (0.006) | 0.306 (0.006) |
|  | CORnet S | 0.643 (0.026) | 0.637 (0.020) | 0.276 (0.027) |
|  | VGG16 | 0.744 (0.006) | 0.707 (0.007) | 0.406 (0.007) |

We sought to assess the behavior of trained models in the same way that we did for Experiment 1. For models trained for single-label classification, we asked whether the models correctly classified images with only a single object annotated. Again, we point out that images with just a single object annotated make up a significant portion of the dataset, as shown in Figure 6. This measure of behavior is no different from that used in Experiment 1, where models assigned images a label of “target present” or “target absent”. Note that when measuring behavior this way with the test set, we used only images where just a single object was annotated. For models trained for multi-label classification, we considered each element in the vector of predictions returned by models as an independent trial, where the model reported that the target was either present or absent. We think this approach is justified, given that our training method for multi-label classification treats the labels that models assign to images as independent. When assessing behavior of models trained this way, we used all images, regardless of the number of objects that were annotated.

To test for a relationship between trial accuracy and difficulty scores, we chose to bin the difficulty scores, then measure trial accuracy of models on the images within each bin. Hence, accuracy within each bin is the number of trials where the model correctly reported a target class was present or absent. We binned the difficulty scores such that there were equal numbers of trials in each bin, so that estimates of accuracy were based on similar sample sizes across bins. For complete details of binning and the trial accuracy calculation, please see training methods and analysis for Visual search difficulty dataset in the Methods. Our analysis revealed a strong correlation between accuracy and difficulty score: as shown in Figure 7, for all models, accuracy dropped as the search difficulty score increased. In both cases, the relationship between accuracy and visual search difficulty score held. We tested different numbers of bins, and found that the effect we saw held regardless. Based on these results, we conclude that DNN models of object recognition exhibit performance limitations when performing a visual search task using complex natural images that is predictive of task difficulty for human participants performing the same task.

As in Experiment 1, we considered the possibility that our results may be an artifact of training. We again measured the accuracy of models on a held-out test set after training, as shown in Table 2. We used the exact same transfer learning method to adapt the object recognition models to both tasks, so that we could directly compare model behavior between tasks. Compared to Experiment 1, models achieved relatively low accuracies on the test set, with ranges from 48-78% when trained for single-label classification on the largest object in each image. We find few reports of single-label image classification accuracy for Pascal VOC, because the dataset usually serves as a benchmark for what are known in machine learning as object detection tasks, where a DNN model learns to predict both a label and bounding box for multiple objects in an image (not to be confused with object recognition tasks in neuroscience). However, we did find one previous report (Shetty, 2016) of 85.6% single-label image classification accuracy for AlexNet pre-trained on ImageNet. In that report the authors fine-tune the weights in all the fully-connected layers, whereas we only train a new last layer. Our training resulted in AlexNet models with 65.2% accuracy. This suggests that simply re-training the last layer as we did likely limits accuracy. Again, we emphasize that our method of retraining just the last layer is necessitated by our goal of asking how object detection models perform visual search tasks. We also emphasize that models with lower accuracy across the entire test set did not necessarily produce higher effect size. We did not observe a strong relationship between accuracy and the R coefficient, as can be seen by comparing Table 2 and Table 3. If anything, models with lower accuracy had less negative R coefficients, suggesting these models showed a weaker inverse correlation with human-derived search difficulty scores. For example, CORnet Z models performed poorly in the multi-label classification setting, achieving the lowest f1 score of 0.113, and that these models had an R coefficient of −0.3, while all other models had an R coefficient of −0.9 or less (by “less” we mean closer to −1.0). Lastly, we recorded and then visually inspected the training histories, as in Experiment 1, and saw in all cases that optimization converged. This can be seen in the training histories for the AlexNet architecture, shown in Figure 8 (plots for other architectures are included in the repository corresponding to this paper). Taken together, these additional results suggest the main effect shown in Figure 7 is not simply due to poor performance of the models.

**Table 3.**

| task (M.L.) | DNN architecture | metric | value | R |
| --- | --- | --- | --- | --- |
| single-label,<br>largest | CORnet Z | mean accuracy<br>(largest object) | 0.54199758 | -0.93413 |
|  | alexnet |  | 0.65150207 | -0.9484 |
|  | CORnet S |  | 0.74305076 | -0.94115 |
|  | VGG16 |  | 0.78578211 | -0.93972 |
| single-label,<br>random | CORnet Z | mean accuracy<br>(random object) | 0.50073377 | -0.9451 |
|  | alexnet |  | 0.58688709 | -0.95905 |
|  | CORnet S |  | 0.65396236 | -0.97148 |
|  | VGG16 |  | 0.70886568 | -0.96891 |
| multi-label | CORnet Z | mean f1<br>(macro) | 0.11255649 | -0.3646 |
|  | alexnet |  | 0.30629583 | -0.93903 |
|  | CORnet S |  | 0.27626449 | -0.9227 |
|  | VGG16 |  | 0.40626116 | -0.97865 |

**Figure 8.**
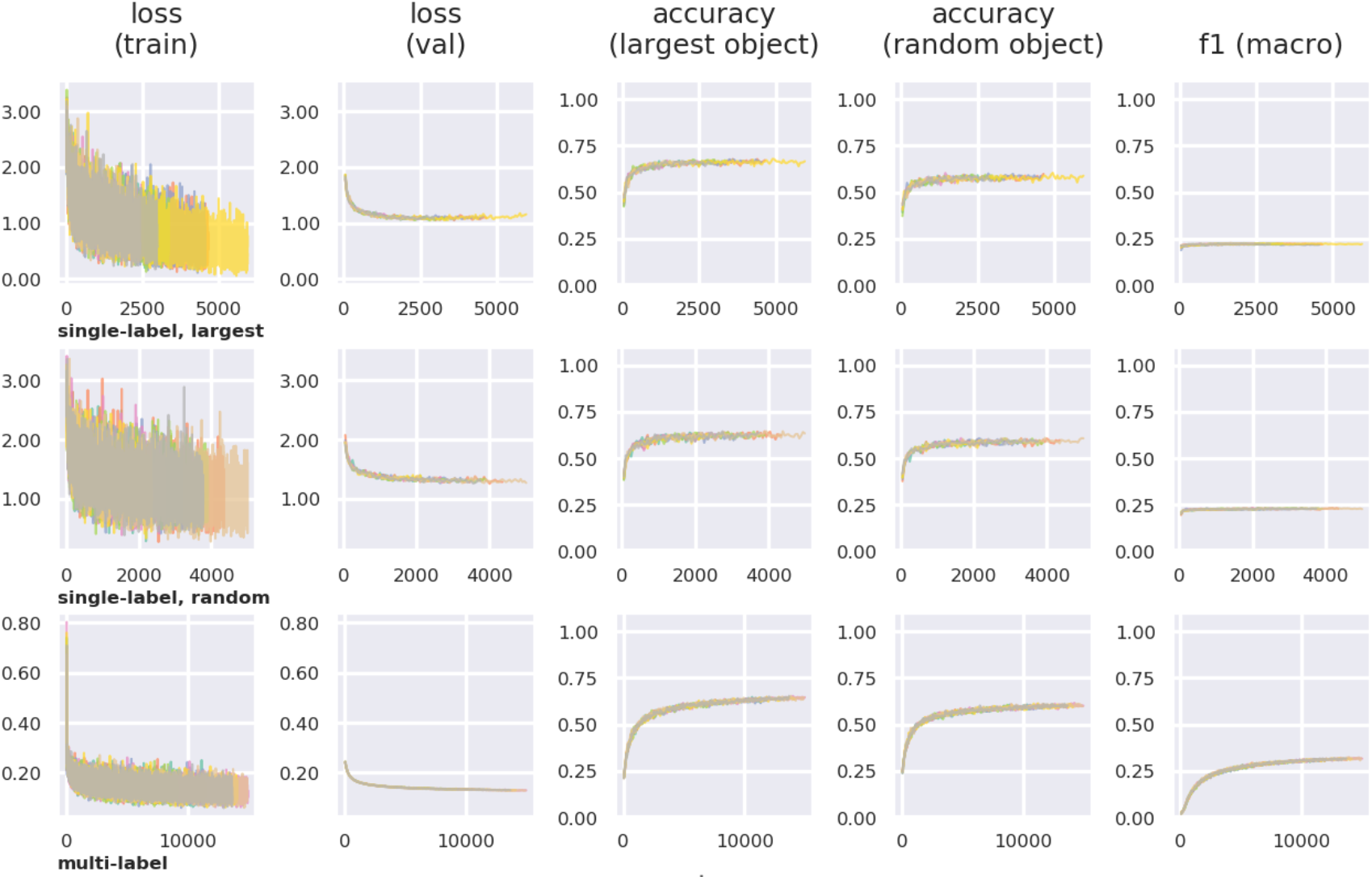
Training histories for DNN architecture trained on visual search task with Pascal VOC dataset. Shown are representative training histories from the Alexnet model. (Plots of training histories for all models can be found in the openly shared repository accompanying this paper; see methods for link.) Each row corresponds to a different machine learning task and/or optimization function as described in text. Hue of lines in each plot indicate training replicates. Each plot shows different metrics measured at different “steps” of training (a “step” is one batch of training data). First column shows the loss function used to optimize the model. Second column shows loss measured intermittently on a validation data set (not used to update model weights). The third through fifth columns show different accuracy metrics, measured on the same validation set.

In Experiment 1, we showed that models trained from randomly-initialized weights were able to achieve near-perfect accuracy when classifying simplified search stimuli as “target present” or “target absent”. One might reasonably ask why not carry out the same experiment here. For this experiment with natural images, we would expect that models trained from randomly-initialized weights would not achieve higher accuracy than models with weights pre-trained for image classification on ImageNet, because transfer learning is known to provide better accuracy than training “from scratch”. In particular this has been shown when pre-training models on ImageNet and then adapting the models to a related task where we have a much smaller training set (Yosinski et al., 2014; Kornblith et al., 2019), as is the case here. For the sake of completeness, we did train models from randomly-initialized weights using single- and multi-label classification, and did find that those models achieved much lower accuracy on the test set. Please see on-line repository for complete results. It could be argued that in this case we are simply training models to perform object recognition, not a visual search task. We agree that the distinction is unclear, and return to this point in the discussion. Because models trained from randomly-initialized weights did not achieve high accuracy on the test set, we did not repeat the analysis correlating their accuracies with visual search difficulty scores. We expect the correlation would be low, given that models with lower accuracy exhibit lower correlations when trained with transfer learning, as we showed above.

## DISCUSSION

We asked if object recognition imposes limits on visual search performance. To test this idea, we made use of DNN-based models of object recognition. A key strength of these models is that they are image-computable, meaning that we might be able to use them to account for performance limitations across all types of images used in visual search tasks. When tested with simplified stimuli typically used in laboratory search tasks, like those shown in Figure 1, the DNN models exhibited set size effects that are hallmarks of this task, as shown in Figure 2 and Figure 4. Similarly, when assayed with real-world objects in natural scenes, like those shown in Figure 6, DNN models show a drop in accuracy inversely correlated with visual search difficulty scores derived from reaction times of human subjects searching the images, as shown in Figure 7. To further test whether the set size effects we saw with simplified stimuli were specific to object recognition models, we carried out separate experiments where we trained the same DNN architectures from randomly-initialized weights, instead of using weights pre-trained for image classification, as is typically done for object recognition models (Schrimpf et al., 2018; D. L. Yamins et al., 2014a). This experiment demonstrated that models trained from randomly-initialized weights are capable of performing the task with near-perfect accuracy, as we show in Figure 2 and Figure 4, and so we conclude that the effect is specific to object recognition models. Taken as a whole, our results are consistent with the idea that object recognition places limitations on visual search performance. Of course, this will need to be further tested by more direct comparisons of DNN model behavior with that of human subjects.

This need to further test empirically points to potential weaknesses of our findings. One weakness relates to the set size effects we saw DNN models exhibit in Experiment 1. Set size effects alone do not provide sufficient support for any mechanism that claims to account for performance limitations (Kristjánsson, 2015; Nakayama & Martini, 2011). For this reason, researchers have turned to multiple measures, such as comparisons between distributions (Wolfe et al., 2010), to arbitrate between proposed mechanisms. It is very possible that measuring multiple aspects of the behavior of the DNN models we tested here may reveal differences in how they solve visual search tasks compared to humans. As others have argued, careful comparisons of the behavior of DNN models and humans are both necessary and informative (Firestone, 2020; Funke et al., 2020; Geirhos et al., 2020; Kim et al., 2020). In spite of any possible differences, we can at least claim that optimizing DNN models for object recognition produces a key effect that has been the focus of many studies of visual search, and in that sense is no weaker than the prevailing models. Another weakness of our approach here is that the DNN models we employed cannot account for other behavioral measures, the most crucial of which is reaction time. We address how to extend models to account for reaction times below, but here we point out that DNN models appear capable of accounting for effects on accuracy seen when participant’s responses are also time-limited. We also showed that DNN accuracy is predictive of a score derived from human reaction times when searching natural images. Hence our results suggest that DNN models have the potential to account for behavior across visual search tasks, a key issue with prevailing models that we set out to address.

A first step towards addressing some of the potential weaknesses of DNN models we have just outlined would be to extend these models so that they also produce reaction times. This would enable researchers to test whether a single model accounts for results not just across stimuli, but also across the different protocols for performing visual search tasks. There are several methods for extending DNN models so they produce reaction times. The first would be to use recurrent neural networks, which carry out a computation for a specified number of time steps *t*, as has been done for studies of object recognition (Kar et al., 2019; Kietzmann et al., 2019; Nayebi et al., 2018; Spoerer et al., 2017). In general, these studies find that recurrence conveys an advantage in terms of predicting neural activity and behavior. Another solution would be to add computations to DNNs from modeling studies of visual search, computations that also produce reaction times, such as a winner-take-all or diffusion-drift mechanisms (Moran et al., 2013; Narbutas et al., 2017). Although these mechanisms could be applied to DNN models, the models would always produce the same reaction time given a particular image, because DNN output is deterministic (at least, at inference time, ignoring things like stochastic dropout often used during training). In contrast, human subjects produce a distribution of reaction times across trials (Wolfe et al., 2010). Addressing all of these factors may require adopting a different theoretical framework. For example, building models within the Neural Engineering Framework (Eliasmith & Anderson, 2003; Eliasmith & Stewart, 2011) would make it possible to combine DNN models tested here (Rasmussen, 2019) with higher-level cognitive mechanisms such as a winner-take-all (Gosmann et al., 2017), while also assessing the role of behavioral stochasticity arising in part from noisy neural activity (Bekolay et al., 2014; Hunsberger et al., 2014; Hunsberger, Eric, 2018).

Another question that might be raised about our approach is whether we have really trained DNNs to perform visual search tasks. Is anything about our methods different from those employed by researchers studying object recognition? We used some of the same benchmark datasets from computer vision, the same models, and the same theoretical framework. Most crucially, we study the exact same behavioral measure, accuracy, and directly compare with human behavior when performing the task in a time-limited setting. Although the tasks researchers use to study these two behaviors are very similar, they are arrived at by different constraints. Researchers studying object recognition have taken this approach in part because they want to compare neural activity in some fixed time window with activation in the DNN models. Usually, activations in the neural network are assumed to be similar to an initial, largely feedforward volley of activity in the ventral visual stream, within the first 100 ms of stimulus onset. (For extensive discussion of this operationalization of object recognition see (Cadieu et al., 2014; Majaj et al., 2015) and references therein.) Researchers studying visual search have also often carried out experiments where they limit the time that subjects view a stimulus, typically to hundreds of milliseconds. In this case, though, the experimental design is motivated by the need to control other factors such as eye movement (Eckstein, 1998, 2011; Wolfe, 1998). In spite of these different constraints, it appears that researchers studying different aspects of vision have arrived at the same “task” for investigating their behavior of interest. The similarity between tasks is most apparent when both researchers use complex natural images as part of their task, as we did in Experiment 2.

Setting aside questions about object recognition and visual search tasks for a moment, we emphasize the clear difference we saw between the two tasks we used here. We showed that DNNs can find near-optimal solutions for the task with simplified stimuli that we used in Experiment 1, when models were trained from randomly-initialized weights, whereas training DNNs the same way for the task in Experiment 2 that uses natural images does not allow them to achieve near-perfect accuracy, and based on results from others (Shetty, 2016), this would be true even if we were not constrained by training data. This clear difference between tasks highlights the questions that remain unanswered within studies of visual search. As others have observed, more work needs to be done to understand whether well-established tasks truly allow us to interrogate how the brain solves problems to produce real-world behaviors (Juavinett et al., 2018; Krakauer et al., 2017). This need to understand what different tasks are telling us is made all the more apparent when we realize that researchers studying two different real-world behaviors—object recognition and visual search—use essentially the same task, as we have just discussed.

Hence our findings principally highlight that prevailing models of visual search neglect object recognition, and further underscore the need to continue study of how the two behaviors are related. How then can future work better clarify this? As stated in the introduction, prevailing models of visual search attribute performance limitations to selective attention mechanisms. Issues with the concept of attention have been widely discussed (Anderson, 2011; Awh et al., 2012; Hommel et al., 2019; Krauzlis et al., 2014; Maunsell, 2004, 2015). In that regard, we point out that “selective attention” is often understood by researchers studying visual search as a source of performance *limitations*, while “attention” is usually understood by researchers in deep learning as some sort of learned weighting that results in performance *improvements*. Interdisciplinary studies reconciling these apparently conflicting definitions of attention might be fruitful (Lindsay, 2020; Marblestone et al., 2016), by using tasks like those we have presented here. In contrast to visual search, researchers studying object recognition assert that it occurs through an untangling mechanism (DiCarlo et al., 2012; DiCarlo & Cox, 2007). DNN models are one implementation of this theoretical mechanism; they achieve “untangling” by performing a series of non-linear transformations on their inputs. As the models’ behavior demonstrates, this series of transformations makes it possible for DNNs to distinguish two objects that may lie very near each other in high-dimensional, low-level feature space. Our results suggest that untangling alone does not limit performance, since we showed that DNNs trained from randomly-initialized weights can find near-perfect solutions to visual search tasks. Instead, we showed that optimizing models for one task limits models’ ability to perform another task. One might argue that the two real-world behaviors are controlled by separate cognitive modules, and it is a mistake to model this by asking a single DNN to perform both tasks. We do not think this is the case: given that in the real world we obviously use our whole brain to produce behavior, it seems worth asking how different behaviors might impose constraints on a given system within the brain. In the worst-case scenario, models of cognition that relegate object recognition and visual search to completely separate modules might be an artifact of psychological history, unconstrained by what we know about how the visual system evolved in concert with the rest of the brain (Cisek, 2019). One way to eliminate this is to push further on DNN models, constrained by results obtained through the study of both object recognition and visual search. Such an approach can force us to build better theories (Guest & Martin, 2020) about how the two behaviors are related in the real world. In this way, the deep learning framework can help answer important questions in neuroscience (Saxe et al., 2020) and related fields even if some of its stronger assertions do not hold.

## CONCLUSIONS

We asked whether object recognition imposes limits on visual search performance across stimuli. To test this idea, we leveraged the strengths of deep neural network (DNN) models of object recognition. We demonstrated that DNNs exhibit a hallmark effect seen when participants search simplified stimuli often used in laboratory tasks, and this effect results from optimizing DNNs for object recognition. Then we showed that with a dataset of relatively complex natural images containing real objects that DNN accuracy is inversely correlated with visual search difficulty scores derived from human reaction times. Taken as a whole, our results are consistent with the idea that object recognition places limitations on visual search performance.

## ACKNOWLEDGEMENTS

Research funded by the Lifelong Learning Machines program, DARPA/Microsystems Technology Office, DARPA cooperative agreement HR0011-18-2-0019. David Nicholson was partially supported by the 2017 William K. and Katherine W. Estes Fund to F. Pestilli, R. Goldstone and L. Smith, Indiana University Bloomington.

## Notes

### Competing Interest Statement

The authors have declared no competing interest.

## REFERENCES

Anderson, B. (2011). There is no such thing as attention. Frontiers in Psychology, 2(SEP), 1–8. https://doi.org/10.3389/fpsyg.2011.00246

Awh, E., Belopolsky, A. V., & Theeuwes, J. (2012). Top-down versus bottom-up attentional control: A failed theoretical dichotomy. Trends in Cognitive Sciences, 16(8), 437–443. https://doi.org/10.1016/j.tics.2012.06.010

Bakdash, J. Z., & Marusich, L. R. (2017). Repeated Measures Correlation. Frontiers in Psychology, 8, 456. https://doi.org/10.3389/fpsyg.2017.00456

Bauer, B., Jolicoeur, P., & Cowan, W. B. (1996). Visual search for colour targets that are or are not linearly separable from distractors. Vision Research, 36(10), 1439–1466.

Bekolay, T., Bergstra, J., Hunsberger, E., DeWolf, T., Stewart, T. C., Rasmussen, D., Choo, X., Voelker, A. R., & Eliasmith, C. (2014). Nengo: A Python tool for building large-scale functional brain models. Frontiers in Neuroinformatics, 7.https://doi.org/10.3389/fninf.2013.00048

Bobier, B., Stewart, T. C., & Eliasmith, C. (2014). A unifying mechanistic model of selective attention in spiking neurons. PLoS Computational Biology, 10(6), e1003577.

Cadieu, C. F., Hong, H., Yamins, D. L. K., Pinto, N., Ardila, D., Solomon, E. A., Majaj, N. J., & DiCarlo, J. J. (2014). Deep Neural Networks Rival the Representation of Primate IT Cortex for Core Visual Object Recognition. PLoS Computational Biology, 10(12), e1003963. https://doi.org/10.1371/journal.pcbi.1003963

Caswell, T. A., Droettboom, M., Lee, A., Hunter, J., Firing, E., Stansby, D., Klymak, J., Hoffmann, T., de Andrade, E. S., Varoquaux, N., Nielsen, J. H., Root, B., Elson, P., May, R., Dale, D., Lee, J.-J., Seppänen, J. K., McDougall, D., Straw, A.,… Katins, J. (2020). Matplotlib/matplotlib v3.1.3 (v3.1.3) [Computer software]. Zenodo. https://doi.org/10.5281/zenodo.3633844

Cisek, P. (2019). Resynthesizing behavior through phylogenetic refinement. Attention, Perception, & Psychophysics, 81(7), 2265–2287. https://doi.org/10.3758/s13414-019-01760-1

Cohen, M. A., Alvarez, G. A., Nakayama, K., & Konkle, T. (2017). Visual search for object categories is predicted by the representational architecture of high-level visual cortex. Journal of Neurophysiology, 117(1), 388–402. https://doi.org/10.1152/jn.00569.2016

Cooper, R. P., & Guest, O. (2014). Implementations are not specifications: Specification, replication and experimentation in computational cognitive modeling. Cognitive Systems Research, 27, 42–49. https://doi.org/10.1016/j.cogsys.2013.05.001

DiCarlo, J. J., & Cox, D. D. (2007). Untangling invariant object recognition. Trends in Cognitive Sciences, 11(8), 333–341.

DiCarlo, J. J., Zoccolan, D., & Rust, N. C. (2012). How Does the Brain Solve Visual Object Recognition? Neuron, 73(3), 415–434. https://doi.org/10.1016/j.neuron.2012.01.010

Duncan, J., & Humphreys, G. W. (1989). Visual Search and Stimulus Similarity. 96(3), 433–458.

D’Zmura, M. (1991). Color in visual search. Vision Research, 31(6), 951–966.

Eckstein, M. P. (1998). The lower visual search efficiency for conjunctions is due to noise and not serial attentional processing. Psychological Science, 9(2), 111–118.

Eckstein, M. P. (2011). Visual search: A retrospective. Journal of Vision, 11(5), 14–14.

Eckstein, M. P., Thomas, J. P., Palmer, J., & Shimozaki, S. S. (2000). A signal detection model predicts the effects of set size on visual search accuracy for feature, conjunction, triple conjunction, and disjunction displays. Perception & Psychophysics, 62(3), 425–451.

Eliasmith, C., & Anderson, C. H. (2003). Neural engineering: Computation, representation, and dynamics in neurobiological systems. MIT press.

Eliasmith, C., & Stewart, T. (2011). Nengo and the neural engineering framework: Connecting cognitive theory to neuroscience. Proceedings of the Annual Meeting of the Cognitive Science Society, 33, Article 33.

Everingham, M., Van Gool, L., Williams, C., Winn, J., & Zisserman, A. (2012). The Pascal visual object classes challenge 2012 results, vol. 5 (2012).

Firestone, C. (2020). Performance vs. Competence in human–machine comparisons. Proceedings of the National Academy of Sciences, 117(43), 26562–26571. https://doi.org/10.1073/pnas.1905334117

Funke, C. M., Borowski, J., Stosio, K., Brendel, W., Wallis, T. S. A., & Bethge, M. (2020). Five Points to Check when Comparing Visual Perception in Humans and Machines. ArXiv:2004.09406 [Cs, q-Bio, Stat]. http://arxiv.org/abs/2004.09406

Geirhos, R., Jacobsen, J.-H., Michaelis, C., Zemel, R., Brendel, W., Bethge, M., & Wichmann, F. A. (2020). Shortcut Learning in Deep Neural Networks. ArXiv:2004.07780 [Cs, q-Bio]. http://arxiv.org/abs/2004.07780

Geisler, W. S. (2003). Ideal observer analysis. The Visual Neurosciences, 10(7), 12–12.

Geisler, W. S., & Cormack, L. K. (2011). Models of overt attention. Oxford Handbook of Eye Movements, 439–454.

Glorot, X., Bordes, A., & Bengio, Y. (2011). Deep sparse rectifier neural networks. Proceedings of the Fourteenth International Conference on Artificial Intelligence and Statistics, 315–323.

Gosmann, J., Voelker, A., & Eliasmith, C. (2017). A spiking independent accumulator model for winner-take-all computation. CogSci.

Green, D. M., Swets, J. A., & others. (1966). Signal detection theory and psychophysics (Vol. 1). Wiley New York.

Grisel, O., Mueller, A., Lars, Gramfort, A., Louppe, G., Prettenhofer, P., Blondel, M., Niculae, V., Nothman, J., Joly, A., Fan, T. J., Vanderplas, J., kumar, manoj, Qin, H., Hug, N., Varoquaux, N., Estève, L., Layton, R., Metzen, J. H.,… du Boisberranger, J. (2020). scikit-learn/scikit-learn: Scikit-learn 0.24.0 (0.24.0) [Computer software]. Zenodo. https://doi.org/10.5281/zenodo.4385486

Grossberg, S., Mingolla, E., & Ross, W. D. (1994). A neural theory of attentive visual search: Interactions of boundary, surface, spatial, and object representations. Psychological Review, 101(3), 470.

Guest, O., & Martin, A. E. (2020). How computational modeling can force theory building in psychological science [Preprint]. PsyArXiv. https://doi.org/10.31234/osf.io/rybh9

Harris, C. R., Millman, K. J., van der Walt, S. J., Gommers, R., Virtanen, P., Cournapeau, D., Wieser, E., Taylor, J., Berg, S., Smith, N. J., & others. (2020). Array programming with NumPy. Nature, 585(7825), 357–362.

Hastie, T., Tibshirani, R., & Friedman, J. (2001). The Elements of Statistical Learning. The Mathematical Intelligencer, 27(2), 83–85. https://doi.org/10.1198/jasa.2004.s339

Hommel, B., Chapman, C. S., Cisek, P., Neyedli, H. F., Song, J.-H., & Welsh, T. N. (2019). No one knows what attention is. Attention, Perception, & Psychophysics, 81(7), 2288–2303. https://doi.org/10.3758/s13414-019-01846-w

Hulleman, J., & Olivers, C. N. (2017a). On the brink: The demise of the item in visual search moves closer. Behavioral and Brain Sciences, 40.

Hulleman, J., & Olivers, C. N. L. (2017b). The impending demise of the item in visual search. Behavioral and Brain Sciences, 40, e132. Cambridge Core. https://doi.org/10.1017/S0140525X15002794

Hunsberger, E., Scott, M., & Eliasmith, C. (2014). The Competing Benefits of Noise and Heterogeneity in Neural Coding. Neural Computation, 26(8), 1600–1623. https://doi.org/10.1162/NECO_a_00621

Hunsberger, Eric. (2018). Spiking Deep Neural Networks: Engineered and Biological Approaches to Object Recognition. UWSpace. http://hdl.handle.net/10012/12819

Hunter, J. D. (2007). Matplotlib: A 2D graphics environment. Computing in Science & Engineering, 9(3), 90–95. https://doi.org/10.1109/MCSE.2007.55

Ionescu, R. T., Alexe, B., Leordeanu, M., Popescu, M., Papadopoulos, D. P., & Ferrari, V. (2016). How Hard Can It Be? Estimating the Difficulty of Visual Search in an Image. 2016 IEEE Conference on Computer Vision and Pattern Recognition (CVPR), 2157–2166. https://doi.org/10.1109/CVPR.2016.237

Jozwik, K. M., Kriegeskorte, N., Storrs, K. R., & Mur, M. (2017). Deep Convolutional Neural Networks Outperform Feature-Based But Not Categorical Models in Explaining Object Similarity Judgments. Frontiers in Psychology, 8, 1726. https://doi.org/10.3389/fpsyg.2017.01726

Juavinett, A. L., Erlich, J. C., & Churchland, A. K. (2018). Decision-making behaviors: Weighing ethology, complexity, and sensorimotor compatibility. Current Opinion in Neurobiology, 49, 42–50.

Kar, K., Kubilius, J., Schmidt, K., Issa, E. B., & DiCarlo, J. J. (2019). Evidence that recurrent circuits are critical to the ventral stream’s execution of core object recognition behavior. Nature Neuroscience, 22(6), 974–983. https://doi.org/10.1038/s41593-019-0392-5

Katti, H., Peelen, M. V., & Arun, S. P. (2017). How do targets, nontargets, and scene context influence real-world object detection? Attention, Perception, & Psychophysics, 79(7), 2021–2036. https://doi.org/10.3758/s13414-017-1359-9

Kell, A. J., & McDermott, J. H. (2019). Deep neural network models of sensory systems: Windows onto the role of task constraints. Current Opinion in Neurobiology, 55, 121–132.

Kietzmann, T. C., Spoerer, C. J., Sörensen, L. K. A., Cichy, R. M., Hauk, O., & Kriegeskorte, N. (2019). Recurrence is required to capture the representational dynamics of the human visual system. Proceedings of the National Academy of Sciences, 116(43), 21854–21863. https://doi.org/10.1073/pnas.1905544116

Kim, B., Reif, E., Wattenberg, M., Bengio, S., & Mozer, M. C. (2020). Neural Networks Trained on Natural Scenes Exhibit Gestalt Closure. ArXiv:1903.01069 [Cs, Stat]. http://arxiv.org/abs/1903.01069

Kluyver, T., Ragan-Kelley, B., Pérez, F., Granger, B., Bussonnier, M., Frederic, J., Kelley, K., Hamrick, J., Grout, J., Corlay, S., Ivanov, P., Avila, D., Abdalla, S., & Willing, C. (2016). Jupyter Notebooks – a publishing format for reproducible computational workflows. In F. Loizides & B. Schmidt (Eds.), Positioning and power in academic publishing: Players, agents and agendas (pp. 87–90).

Kornblith, S., Shlens, J., & Le, Q. V. (2019). Do better imagenet models transfer better? Proceedings of the IEEE Conference on Computer Vision and Pattern Recognition, 2661–2671.

Krakauer, J. W., Ghazanfar, A. A., Gomez-Marin, A., MacIver, M. A., & Poeppel, D. (2017). Neuroscience needs behavior: Correcting a reductionist bias. Neuron, 93(3), 480–490.

Krauzlis, R. J., Bollimunta, A., Arcizet, F., & Wang, L. (2014). Attention as an effect not a cause. Trends in Cognitive Sciences, 18(9), 457–464. https://doi.org/10.1016/j.tics.2014.05.008

Kristjánsson, A. (2015). Reconsidering visual search. I-Perception, 6(6), 2041669515614670.

Krizhevsky, A., Sutskever, I., & Hinton, G. E. (2012). Imagenet classification with deep convolutional neural networks. Advances in Neural Information Processing Systems, 1097–1105.

Li, Y., Song, Y., & Luo, J. (2017). Improving Pairwise Ranking for Multi-label Image Classification. 2017 IEEE Conference on Computer Vision and Pattern Recognition (CVPR), 1837–1845. https://doi.org/10.1109/CVPR.2017.199

Lindsay, G. W. (2020). Attention in Psychology, Neuroscience, and Machine Learning. Frontiers in Computational Neuroscience, 14, 29. https://doi.org/10.3389/fncom.2020.00029

Lindsay, G. W., & Miller, K. (2017). Understanding Biological Visual Attention Using Convolutional Neural Networks. https://doi.org/10.1101/233338

Ma, W. J., Navalpakkam, V., Beck, J. M., Berg, R. van den, & Pouget, A. (2011). Behavior and neural basis of near-optimal visual search. Nature Neuroscience, 14(6), 783–790. https://doi.org/10.1038/nn.2814

Majaj, N. J., Hong, H., Solomon, E. A., & DiCarlo, J. J. (2015). Simple Learned Weighted Sums of Inferior Temporal Neuronal Firing Rates Accurately Predict Human Core Object Recognition Performance. Journal of Neuroscience, 35(39), 13402–13418. https://doi.org/10.1523/JNEUROSCI.5181-14.2015

Marblestone, A., Wayne, G., & Kording, K. (2016). Towards an integration of deep learning and neuroscience. 10(September), 1–41. https://doi.org/10.3389/fncom.2016.00094

Marcel, S., & Rodriguez, Y. (2010). Torchvision the machine-vision package of torch. Proceedings of the 18th ACM International Conference on Multimedia, 1485–1488.

Maunsell, J. H. R. (2004). Neuronal representations of cognitive state: Reward or attention? Trends in Cognitive Sciences, 8(6), 261–265. https://doi.org/10.1016/j.tics.2004.04.003

Maunsell, J. H. R. (2015). Neuronal Mechanisms of Visual Attention. Annual Review of Vision Science, 1(1), 373–391. https://doi.org/10.1146/annurev-vision-082114-035431

Moran, R., Zehetleitner, M., Muller, H. J., & Usher, M. (2013). Competitive guided search: Meeting the challenge of benchmark RT distributions. Journal of Vision, 13(8), 24–24. https://doi.org/10.1167/13.8.24

Nakayama, K., & Martini, P. (2011). Situating visual search. Vision Research, 51(13), 1526–1537. https://doi.org/10.1016/j.visres.2010.09.003

Nam, J., Kim, J., Loza Mencía, E., Gurevych, I., & Fürnkranz, J. (2014). Large-Scale Multi-label Text Classification—Revisiting Neural Networks. In T. Calders, F. Esposito, E. Hüllermeier, & R. Meo (Eds.), Machine Learning and Knowledge Discovery in Databases (Vol. 8725, pp. 437–452). Springer Berlin Heidelberg. https://doi.org/10.1007/978-3-662-44851-9_28

Narbutas, V., Lin, Y.-S., Kristan, M., & Heinke, D. (2017). Serial versus parallel search: A model comparison approach based on reaction time distributions. Visual Cognition, 25(1–3), 306–325. https://doi.org/10.1080/13506285.2017.1352055

Nayebi, A., Bear, D., Kubilius, J., Kar, K., Ganguli, S., Sussillo, D., DiCarlo, J. J., & Yamins, D. L. (2018). Task-Driven Convolutional Recurrent Models of the Visual System. ArXiv Preprint ArXiv:1807.00053.

Nicholson, D., & Prinz, A. (2019). Convolutional neural networks performing a visual search task show attention-like limits on accuracy when trained to generalize across multiple search stimuli. 2019 Conference on Cognitive Computational Neuroscience. 2019 Conference on Cognitive Computational Neuroscience, Berlin, Germany. https://doi.org/10.32470/CCN.2019.1432-0

Palmer, E. M., Fencsik, D. E., Flusberg, S. J., Horowitz, T. S., & Wolfe, J. M. (2011). Signal detection evidence for limited capacity in visual search. Attention, Perception, & Psychophysics, 73(8), 2413–2424.

Palmer, J. (1994). Set-size effects in visual search: The effect of attention is independent of the stimulus for simple tasks. Vision Research, 34(13), 1703–1721.

Palmer, J., Verghese, P., & Pavel, M. (2000). The psychophysics of visual search. Vision Research, 40(10–12), 1227–1268.

Paszke, A., Gross, S., Massa, F., Lerer, A., Bradbury, J., Chanan, G., Killeen, T., Lin, Z., Gimelshein, N., Antiga, L., Desmaison, A., Kopf, A., Yang, E., DeVito, Z., Raison, M., Tejani, A., Chilamkurthy, S., Steiner, B., Fang, L.,… Chintala, S. (2019). PyTorch: An imperative style, high-performance deep learning library. In H. Wallach, H. Larochelle, A. Beygelzimer, F. dAlché-Buc, E. Fox, & R. Garnett (Eds.), Advances in neural information processing systems 32 (pp. 8024–8035). Curran Associates, Inc. http://papers.neurips.cc/paper/9015-pytorch-an-imperative-style-high-performance-deep-learning-library.pdf

Pedregosa, F., Varoquaux, G., Gramfort, A., Michel, V., Thirion, B., Grisel, O., Blondel, M., Prettenhofer, P., Weiss, R., Dubourg, V., Vanderplas, J., Passos, A., Cournapeau, D., Brucher, M., Perrot, M., & Duchesnay, E. (2011). Scikit-learn: Machine learning in Python. Journal of Machine Learning Research, 12, 2825–2830.

Peelen, M. V., & Kastner, S. (2014). Attention in the real world: Toward understanding its neural basis. Trends in Cognitive Sciences, 18(5), 242–250.

Poder, E. (2017). Capacity limitations of visual search in deep convolutional neural network. ArXiv Preprint ArXiv:1707.09775.

Rasmussen, D. (2019). NengoDL: Combining deep learning and neuromorphic modelling methods. ArXiv:1805.11144 [Cs]. http://arxiv.org/abs/1805.11144

Richards, B. A., Lillicrap, T. P., Beaudoin, P., Bengio, Y., Bogacz, R., Christensen, A., Clopath, C., Costa, R. P., de Berker, A., & Ganguli, S. (2019). A deep learning framework for neuroscience. Nature Neuroscience, 22(11), 1761–1770.

Saxe, A., Nelli, S., & Summerfield, C. (2020). If deep learning is the answer, what is the question? Nature Reviews Neuroscience. https://doi.org/10.1038/s41583-020-00395-8

Schinners, P. (2019). PyGame 1.9.6.

Schlawack, H. (2019). Attrs 19.3.0.

Schrimpf, M., Kubilius, J., Hong, H., Majaj, N. J., Rajalingham, R., Issa, E. B., Kar, K., Bashivan, P., Prescott-Roy, J., & Schmidt, K. (2018). Brain-Score: Which artificial neural network for object recognition is most brainlike? BioRxiv, 407007.

Seabold, S., & Perktold, J. (2010). statsmodels: Econometric and statistical modeling with python. 9th Python in Science Conference.

Shetty, S. (2016). Application of convolutional neural network for image classification on Pascal VOC challenge 2012 dataset. ArXiv Preprint ArXiv:1607.03785.

Simonyan, K., & Zisserman, A. (2014). Very deep convolutional networks for large-scale image recognition. ArXiv Preprint ArXiv:1409.1556.

Spoerer, C. J., McClure, P., & Kriegeskorte, N. (2017). Recurrent Convolutional Neural Networks: A Better Model of Biological Object Recognition. Frontiers in Psychology, 8, 1551. https://doi.org/10.3389/fpsyg.2017.01551

team, T. pandas development. (2020). pandas-dev/pandas: Pandas (latest) [Computer software]. Zenodo. https://doi.org/10.5281/zenodo.3509134

Treisman, A. M., & Gelade, G. (1980). A feature-integration theory of attention. Cognitive Psychology, 12(1), 97136.

Vallat, R. (2018). Pingouin: Statistics in Python. Journal of Open Source Software, 3(31), 1026. https://doi.org/10.21105/joss.01026

Virtanen, P., Gommers, R., Oliphant, T. E., Haberland, M., Reddy, T., Cournapeau, D., Burovski, E., Peterson, P., Weckesser, W., Bright, J., van der Walt, S. J., Brett, M., Wilson, J., Millman, K. J., Mayorov, N., Nelson, A. R. J., Jones, E., Kern, R., Larson, E.,… Contributors, S. 1 0. (2019). SciPy 1.0–Fundamental Algorithms for Scientific Computing in Python. ArXiv:1907.10121 [Physics]. http://arxiv.org/abs/1907.10121

Walt, S. van der, Colbert, S. C., & Varoquaux, G. (2011). The NumPy Array: A Structure for Efficient Numerical Computation. Computing in Science Engineering, 13(2), 22–30. https://doi.org/10.1109/MCSE.2011.37

Wang, J., Yang, Y., Mao, J., Huang, Z., Huang, C., & Xu, W. (2016). CNN-RNN: A Unified Framework for Multi-label Image Classification. 2016 IEEE Conference on Computer Vision and Pattern Recognition (CVPR), 2285–2294. https://doi.org/10.1109/CVPR.2016.251

Waskom, M., Botvinnik, O., Ostblom, J., Lukauskas, S., Hobson, P., MaozGelbart, Gemperline, D. C., Augspurger, T., Halchenko, Y., Cole, J. B., Warmenhoven, J., de Ruiter, J., Pye, C., Hoyer, S., Vanderplas, J., Villalba, S., Kunter, G., Quintero, E., Bachant, P.,… Evans, C. (2020). mwaskom/seaborn: V0.10.0 (January 2020) (v0.10.0) [Computer software]. Zenodo. https://doi.org/10.5281/zenodo.3629446

Wes McKinney. (2010). Data Structures for Statistical Computing in Python. In S. van der Walt & Jarrod Millman (Eds.), Proceedings of the 9th Python in Science Conference (pp. 56–61). https://doi.org/10.25080/Majora-92bf1922-00a

Wolfe, J. M. (n.d.). Guided Search 4.0. 22.

Wolfe, J. M. (1994). Guided search 2.0 a revised model of visual search. Psychonomic Bulletin & Review, 1(2), 202–238.

Wolfe, J. M. (1998). Visual search. In Attention (pp. 13–73). Psychology Press/Erlbaum (UK) Taylor & Francis.

Wolfe, J. M., Alvarez, G. A., Rosenholtz, R., Kuzmova, Y. I., & Sherman, A. M. (2011). Visual search for arbitrary objects in real scenes. Attention, Perception, & Psychophysics, 73(6), 1650–1671. https://doi.org/10.3758/s13414-011-0153-3

Wolfe, J. M., & Bennett, S. C. (1997). Preattentive Object Files: Shapeless Bundles of Basic Features. Vision Research, 37(1), 25–43. https://doi.org/10.1016/S0042-6989(96)00111-3

Wolfe, J. M., Cave, K. R., & Franzel, S. L. (1989). Guided search: An alternative to the feature integration model for visual search. Journal of Experimental Psychology: Human Perception and Performance, 15(3), 419.

Wolfe, J. M., & Gray, W. (2007). Guided search 4.0. Integrated Models of Cognitive Systems, 99–119.

Wolfe, J. M., & Horowitz, T. S. (2017). Five factors that guide attention in visual search. Nature Human Behaviour, 1(3), 1–8. https://doi.org/10.1038/s41562-017-0058

Wolfe, J. M., Palmer, E. M., & Horowitz, T. S. (2010). Reaction time distributions constrain models of visual search. Vision Research, 50(14), 1304–1311.

Yamins, D. L., Hong, H., Cadieu, C. F., Solomon, E. A., Seibert, D., & DiCarlo, J. J. (2014a). Performance-optimized hierarchical models predict neural responses in higher visual cortex. Proceedings of the National Academy of Sciences, 111(23), 8619–8624.

Yamins, D. L., Hong, H., Cadieu, C. F., Solomon, E. A., Seibert, D., & DiCarlo, J. J. (2014b). Performance-optimized hierarchical models predict neural responses in higher visual cortex. Proceedings of the National Academy of Sciences, 111(23), 8619–8624.

Yamins, D. L. K., & DiCarlo, J. J. (2016). Using goal-driven deep learning models to understand sensory cortex. Nature Neuroscience, 19(3), 356–365. https://doi.org/10.1038/nn.4244

Yosinski, J., Clune, J., Bengio, Y., & Lipson, H. (2014). How transferable are features in deep neural networks? Advances in Neural Information Processing Systems, 3320–3328.

